# The mitogenome of Norway spruce and a reappraisal of mitochondrial recombination in plants

**DOI:** 10.1101/682732

**Authors:** Alexis R. Sullivan, Yrin Eldfjell, Bastian Schiffthaler, Nicolas Delhomme, Torben Asp, Kim H. Hebelstrup, Olivier Keech, Lisa Öberg, Ian Max Møller, Lars Arvestad, Nathaniel R. Street, Xiao-Ru Wang

## Abstract

Plant mitogenomes can be difficult to assemble because they are structurally dynamic and prone to intergenomic DNA transfers, leading to the unusual situation where an organelle genome is far outnumbered by its nuclear counterparts. As a result, comparative mitogenome studies are in their infancy and some key aspects of genome evolution are still known mainly from pre-genome, qualitative methods. To help address these limitations, we combined machine learning and *in silico* enrichment of mitochondrial-like long reads to assemble the bacterial-sized mitogenome of Norway spruce (Pinaceae: *Picea abies*). We conducted comparative analyses of repeat abundance, intergenomic transfers, substitution and rearrangement rates, and estimated repeat-by-repeat homologous recombination rates. Prompted by our discovery of highly recombinogenic small repeats in *P. abies*, we assessed the genomic support for the prevailing hypothesis that intramolecular recombination is predominantly driven by repeat length, with larger repeats facilitating DNA exchange more readily. Overall, we found mixed support for this view: recombination dynamics were heterogeneous across vascular plants and highly active small repeats (*ca*. 200 bp) were present in about a third of studied mitogenomes. As in previous studies, we did not observe any robust relationships among commonly-studied genome attributes, but we identify variation in recombination rates as a underinvestigated source of plant mitogenome diversity.

## Introduction

Mitochondria share an α-proteobacterium ancestor, a conserved core proteome, and an almost universal function as the site of cellular energy production (Gray 2014). Despite the broad similarity of mitochondria across eukaryotes, the vestigial mitogenome is remarkably diverse in size, content, and architecture (Burger et al. 2003). Nowhere is this heterogeneity showcased more clearly than in plants: closely-related mitogenomes can vary 100-fold or more in size (Sloan et al. 2012), substitution rates (Cho et al. 2004), and rearrangement rates (Cole et al. 2018). Gene repertoires are fluid due to recurrent horizontal (Rice et al. 2013; Sanchez-Puerta 2014) and intergenomic transfers (Adams & Palmer 2003). Multichromosomal architectures (Alverson, Rice, et al. 2011; Sloan et al. 2012) have also been reported from the relatively few sequenced plant mitogenomes. Underlying the dynamism of plant mitogenomes appears to be the evolution of pervasive homologous recombination (Palmer & Herbon 1988; Gray et al. 1999).

Recombination is the predominant mode of double-strand break repair in plant mitogenomes (Maréchal & Brisson 2010; Gualberto & Newton 2017). In contrast, animal mitogenomes tend to use non-homologous end joining pathways or may simply degrade damaged molecules (Alexeyev et al. 2013). Recombination can preserve sequence identity, but reliance on this pathway may lead to unsuppressed intramolecular recombination at dispersed repeats (Maréchal & Brisson 2010). While these differences in repair pathways likely contribute to the broad mitogenome differences among eukaryotic kingdoms, the importance of recombination and other mechanisms in generating the diversity within plants is unclear. Our limited understanding stems, in part, from the few available assembled mitogenomes.

While sequencing plant genomes is routine, only one mitogenome is published for every four nuclear genomes (NCBI Genome Resource; accessed March 11, 2019). Often, mitogenome assembly requires isolating DNA from intact mitochondria because frequent intergenomic transfers (e.g. Alverson, Rice, et al. 2011), the rapid decay of intergenic sequence homology (Guo et al. 2016), and the unclear physical organization of the mitogenome (Sloan 2013) can preclude identifying mitochondrial sequences from whole genome read data (e.g. Nystedt et al. 2013). Strategies to identify mitogenomic scaffolds using GC-content and genome copy number can be effective (e.g. Naito et al. 2013), but this simple strategy becomes prone to false positives and low recovery rates with increasingly large, complex genomes (Eldfjell 2018).

We used machine learning to assemble the bacterial-sized mitogenome of a coniferous forest tree, Norway spruce (Pinaceae: *Picea abies*), from whole-genome shotgun sequencing reads. The *P. abies* mitogenome helps to fill a phylogenetic gap in comparative analyses, and to that end we analyzed gene repertoires; sources of genome size heterogeneity; intraspecific variation; and mutation, recombination, and rearrangement rates in gymnosperms. Prompted by the detection of highly recombinogenic small repeats in *P. abies*, we reevaluated published recombination rates in seed plants. Despite early recognition of recombination as a factor in generating the diversity of eukaryotic mitogenomes (Palmer & Herbon 1988; Gray et al. 1999), surprisingly little attention has been given to recombinational dynamics as a source of mitogenomic diversity within plants. As in previous studies, we found no clear relationship between mitogenome traits and potential mechanisms in – for example, genome size and the proportion of intergenomically transferred DNA – but the role of recombination as a driver of mitogenomic diversity within plants merits further scrutiny.

## Results and Discussion

### Mitogenome assembly

We combined machine learning, *in silico* enrichment of long sequencing reads, and assembly reconciliation to produce a highly contiguous *P. abies* draft mitogenome (Fig. 1). First, we developed a support vector machine (SVM) to identify mitochondrial-like scaffolds from the *P. abies* v. 1.0 genome assembly (Nystedt et al. 2013). Training the SVM classifier requires positive and negative training data and a set of variables – that is, genomic features – expected to discriminate between mitochondrial scaffolds and those from other genomes (Fig. 1). In brief, we used the whole genomes of *Arabidopsis thaliana* and *Populus trichocarpa* and 18 organelle genomes as training data and calculated depth of coverage, guanine and cytosine (%GC) content, and a kmer score based on the frequency of nonredundant ‘mitochondrial-like’ oligonucleotide sequences in each scaffold for use as genomic features (see Methods).

**Figure 1.**
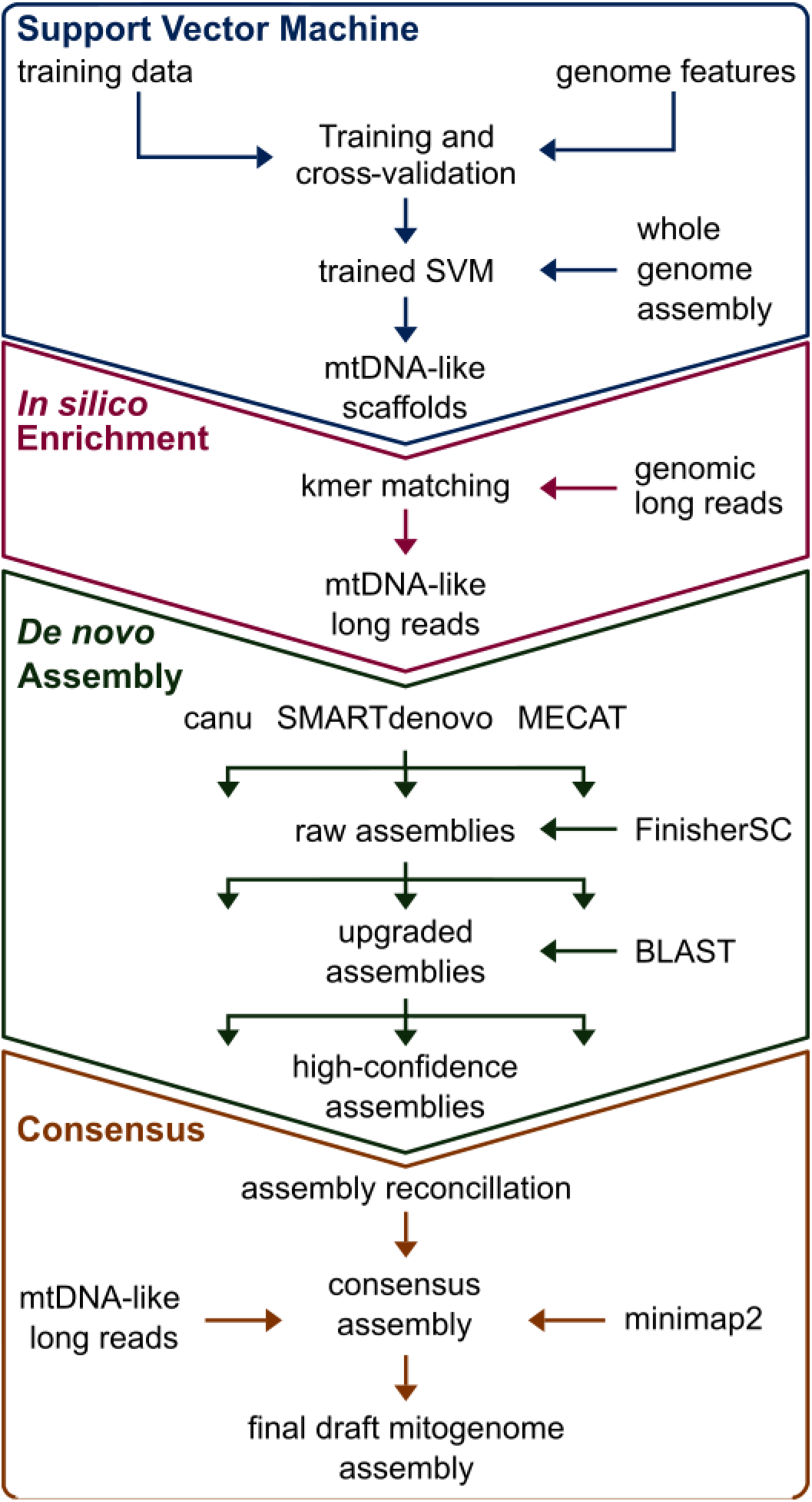
Strategy used to assemble the mitogenome of *Picea abies*. First, a support vector machine (SVM) was trained to identify mitochondrial-like scaffolds from the *P. abies* genome assembly. We used the classified scaffolds to identify PacBio Sequel^®^ subreads containing mitogenome-like 27-mers. These enriched reads were then assembled using three different pipelines. Scaffolds from each assembler with at least one mitochondrial protein coding gene were retained for assembly reconciliation and base-pair correction, thus yielding the final mitogenome draft.

After training the SVM, we applied it to a reduced set of the *P. abies* v. 1.0 assembly (Nystedt et al. 2013) comprising 49,500 scaffolds (see Methods). Of these, the SVM identified 301 scaffolds totaling 4.69 Mb as potentially mitochondrial in origin. While the SVM achieved higher resolution than simple coverage *vs*. GC-content plots (Eldfjell 2018), the recall and false discovery rate (FDR) indicated this classification alone would be insufficient to identify the mitogenome from the *P. abies* v. 1.0 assembly. Recall ranged from 0.53 to 0.92 and the FDR from 0.13 to 0.22 with varying test-to-training ratios (Table S1), although SVMs achieved more accurate classifications in our tests with other draft gymnosperm assemblies (Eldfjell 2018).

We then used the SVM classified scaffolds as ‘bait’ to enrich genomic Pacific Biosciences (PacBio) Sequel^®^ reads for mitochondrial-like sequences *in silico* through kmer matching (Fig. 1). We screened *ca*. 35.5 million subreads totaling 244 Gbp for 27-mer matches to any of the SVM-classified scaffolds, which yielded 3.23 reads totaling 4.1 Gb, with an average read length of 12,041 bp. We reasoned that the long read length could enable us to recover a greater proportion of the mitogenome, as only a single 27-mer match is required, while the improved contiguity of the assembly could allow removal of dubious contigs post-assembly.

We assembled the enriched PacBio reads using three assemblers selected based on their prior performance (Jayakumar & Sakakibara 2017; Giordano et al. 2017; Fig. 1). The most contiguous results per assembler are summarized in Table 1. Upgrading the initial assemblies with unused overlap information in the corrected PacBio reads with FinisherSC (Lam et al. 2015) only appreciably improved the N50 of the canu (Koren et al. 2017) assembly (Table 1). Despite initial differences in size and contiguity, the assemblies were similar when considering only contigs containing at least one of the 41 mitochondrial protein-coding genes conserved in *Gingko* and *Cycas*, which we term high-confidence assemblies (Table 1). Finally, reconciliation using the Contig Integrator for Sequence Assembly (CISA) pipeline (Lin & Liao 2013) produced a final draft assembly measuring 4.9 Mb over four contigs with a mean depth of coverage of 284x.

**Table 1.**
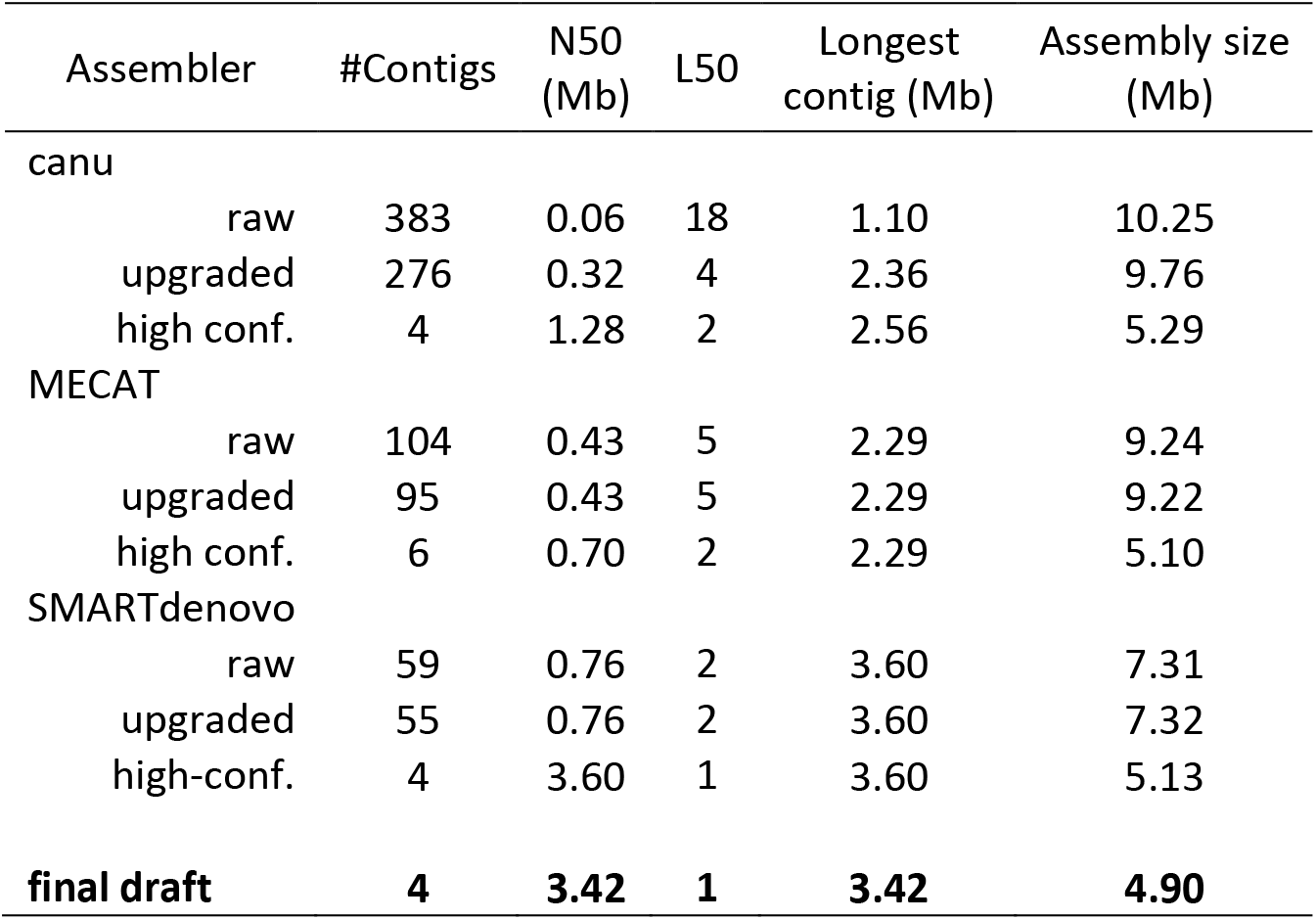
Summary statistics for assemblies of Pacific Biosciences Sequel^®^ subreads enriched *in silico* for mitogenome-like kmers. ‘Raw’ refers to the full contig output produced by each assembler. Upgraded assemblies have been processed with FinisherSC. High-confidence assemblies contain only contigs with at least one protein-coding mitochondrial gene. N50 and L50 are calculated from contig lengths.

| Assembler | #Contigs | N50 (Mb) | L50 | Longest contig (Mb) | Assembly size (Mb) |
| --- | --- | --- | --- | --- | --- |
| canu |  |  |  |  |  |
| raw | 383 | 0.06 | 18 | 1.10 | 10.25 |
| upgraded | 276 | 0.32 | 4 | 2.36 | 9.76 |
| high conf. | 4 | 1.28 | 2 | 2.56 | 5.29 |
| MECAT |  |  |  |  |  |
| raw | 104 | 0.43 | 5 | 2.29 | 9.24 |
| upgraded | 95 | 0.43 | 5 | 2.29 | 9.22 |
| high conf. | 6 | 0.70 | 2 | 2.29 | 5.10 |
| SMARTdenovo |  |  |  |  |  |
| raw | 59 | 0.76 | 2 | 3.60 | 7.31 |
| upgraded | 55 | 0.76 | 2 | 3.60 | 7.32 |
| high-conf. | 4 | 3.60 | 1 | 3.60 | 5.13 |
| **final draft** | **4** | **3.42** | **1** | **3.42** | **4.90** |

### Genome annotation and gene repertoires in gymnosperms

General features of the *P. abies* mitogenome are summarized in Table 2. All 41 prote-incoding genes inferred to be present in the last common ancestor of angiosperms were also found in *P. abies* and, after re-annotation, also in *P. glauca* and *Pinus taeda*. This pattern of conservation in Pinaceae is consistent with the early-diverging gymnosperms *Gingko* and *Cycas*, which may suggest a less dynamic gene repertoire than in angiosperms, although *Welwitschia* has undergone extensive gene loss (Guo et al. 2016). In addition, the *P. abies* mitogenome has also acquired complete copies of four plastid genes, *psaB*, *psbH*, *psbN*, and *psbT*, although the functionality of these genes, if any, cannot be inferred from the data here. We also identified 20 transcribed open-reading frames: 14 had uniquely mapping transcripts of ≥ 99% coverage and identity (high confidence), two were of medium confidence, where either identity or coverage was < 99% but > 97%, and four low confidence genes had less than 97% support (Table S2). Intron and tRNA content were similar in *P. abies* and *P. glauca* (Jackman et al. 2015), but both have apparently undergone losses compared to the other gymnosperms, including the Pinaceae conifer *Pinus taeda*.

**Table 2.**
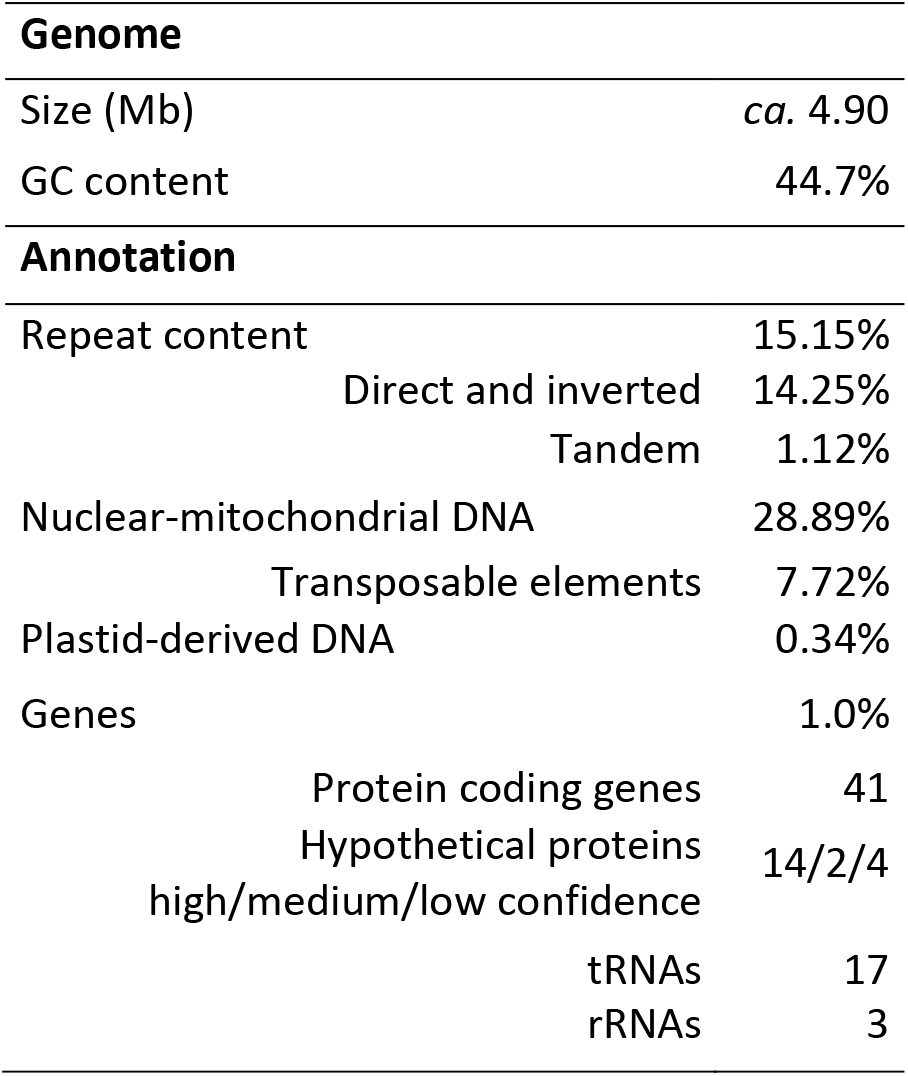
Characteristics of the *Picea abies* mitogenome.

The repetitive fraction of the *P. abies* mitogenome is larger than the entire mitogenome of most plants (Table 2; Table S3). Relative to genome size, however, repeat content in *P. abies* is unremarkable. Dispersed repeats in an analysis of 82 angiosperms comprised 14% of the mitogenome on average (Dong et al. 2018), identical to the proportion in *P. abies* (Table 2). However, dispersed repeats in *P. abies* tended to be about half the size of a typical tracheophyte, at 312 bp *vs*. the 641 bp average (Wynn & Christensen 2019), indicating a relative enrichment of smaller repeats (Fig. S1). At the same time, *P. abies* also had eight pairs of repeats ≥ 10 Kb, more than all but two of the 79 land plants analyzed by Wynn & Christensen (2019).

### Repeats and DNA transfers do not explain *Picea* mitogenome expansion

Extraordinary mitogenome size heterogeneity among plants has long been recognized (Ward et al. 1981), but the sources appear to vary among species. We analyzed intergenomic transfers and the proliferation of repetitive sequences as potential causes of genome size variation in gymnosperms. Transfer of plastid DNA made a negligible contribution to the genome size of the three Pinaceae conifers (Table 3), in contrast to some angiosperms, such as cucurbits (Alverson, Rice, et al. 2011; Alverson et al. 2010). Similarly, no relationship between genome size and repeat proliferation was evident: the 410 kb *Cycas* mitogenome was proportionally the most repetitive (Chaw et al. 2008), and relative repeat content was similar in *Picea* and *Ginkgo* despite their > 10x size differences (Table 3). An ambiguous correlation between genome size and repeat proliferation was also observed in *Silene* species (Sloan et al. 2012), whereas larger cucurbit genomes tend to contain proportionally more repeats (Alverson et al. 2010; Alverson, Rice, et al. 2011; Rodríguez-Moreno et al. 2011).

**Table 3.** Potential sources of mitogenome size variation among gymnosperms. Dispersed repeats include those in the inverted and direct orientation ≥ 50 bp and with ≥ 80% identity. Nuclear-mitochondrial DNA are shared sequences with no direction of transfer inferred. Transposable elements comprises long-terminal repeats (LTR) and non-LTR retrotransposons. Numbers in parenthesis indicate percent coverage of the mitogenome.

|  | *Cycas* | *Ginkgo* | *Picea abies* | *Picea glauca* | *Pinus taeda* | *Welwitschia* |
| --- | --- | --- | --- | --- | --- | --- |
| Genome size (Mb) | 0.41 | 0.35 | 4.90 | 5.20<sup>†</sup> | 1.19 | 0.98 |
| Plastid-derived DNA (kb) | 18 (4) | 0 (0) | 17 (0) | 18 (0) | 3 (0) | 9 (9) |
| Dispersed repeats (kb) | 109 (26) | 51 (15) | 699 (14) | 885 (17) | 83 (7) | 42 (4) |
| Nuclear-mtDNA (Mb) | - | 0.35 (100) | 1.40 (29) | 2.29 (44) | 0.60 (50) | - |
| Transposable elements (kb) | 6 (1) | 3 (1) | 482 (10) | 386 (7) | 129 (10) | 15 (2) |

<sup>†</sup>Gene-containing scaffolds extracted from the putative 5.9 Mb assembly.

Interpreting shared nuclear-mitochondrial content is difficult because 1) the nuclear genomes of *Cycas* and *Welwitschia* are not sequenced; 2) the direction of transfer can rarely be determined with confidence under the simplest circumstances (Alverson et al. 2010); and 3) the *Cycas* and *Ginkgo* mitogenomes are highly conserved, which implies that any import from the nucleus predate their divergence *ca*. 354 My ago (Lu et al. 2014) and may no longer be detectable in either nuclear genome (Guo et al. 2016). Therefore, it is unsurprising that the four species with nuclear reference genomes show an equivocal relationship between mitogenome size and shared nuclear-mitochondrial sequence (Table 3). Remnants of nuclear transposable elements (TEs) can potentially act as markers of sequence import because they have an unambiguous origin and generally do not proliferate after transfer (Knoop et al. 1996; Goremykin et al. 2012). All three Pinaceae species showed an increase of TE remnants, but relative content was similar among the two *Picea* species and the 4-fold smaller *P. taeda* mitogenome (Table 3). Similarly, *Welwitschia* was relatively depauperate in TE elements, despite its large mitogenome size (Table 3). Increased taxon sampling may help to clarify this ambiguous relationship between TE import and genome size, and but a similarly unclear relationship has also been reported among angiosperms (Alverson et al. 2010; Alverson, Rice, et al. 2011; Rodríguez-Moreno et al. 2011, Goremykin et al. 2012). Overall, gymnosperms tend to reinforce observations in angiosperms: repeat proliferation and DNA imports do not broadly explain mitogenome size heterogeneity (Alverson, Rice, et al. 2011; Sloan et al. 2012; Dong et al. 2018; Alverson et al. 2010).

### Abundant repeat-mediated recombination at small repeats

Homologous intermolecular recombination is used to repair double-stranded breaks and recover stalled replication forks (Maréchal & Brisson 2010). In many plants, intramolecular recombination also occurs at dispersed repeats and results in an individual harboring alternative genome configurations (AGCs) that differ in structure but are identical in sequence. Intramolecular recombination dynamics range from completely inert (Alverson, Rice, et al. 2011; Dong et al. 2018) to astonishingly friable (Skippington et al. 2015), yet recombination rates are not widely estimated (see next section). However, recombination resulting in DNA exchange produces predictable AGCs that are directly observable in a pool of single molecule sequencing reads (e.g. Dong et al. 2018; Wang et al. 2018; Fig. S2). This allowed us to identify and estimate the AGC frequency associated with each pair of inverted repeats in the *P. abies* mitogenome, which provides a quantitative estimate of recombination rates.

Genome rearrangements consistent with the products of intramolecular recombination (i.e., AGCs) comprised *ca*. 1% of the pool of mapped reads. Half of the 598 repeats showed no evidence of recombination, but AGC frequency averaged 2% at recombinogenic repeats (Fig. 2A; Table S4). Recombination was asymmetric in most cases, a result consistent with the repair of stalled replication forks through break-induced replication (Table S4; Maréchal & Brisson 2010). AGC abundance reached a maximum of 32% at a 186 bp repeat, with the two possible isomers comprising 12% and 20% of the reads, respectively. Surprisingly, the most recombinogenic repeats (AGCs ≥ 10%) ranged in size from 50 bp, the minimum size evaluated, to 948 bp (Fig. 2A; Table S4). Overall, we found negligible correlation between repeat length and AGC frequency (Fig. 2B; r^2^ = 0.03, p < 0.001), in contrast to some well-characterized examples (e.g. Mower, Floro, et al. 2012; Skippington et al. 2015; Guo et al. 2016) and contrary to the general expectation that, in the absence of other factors, recombination should scale positively with repeat length (Arrieta-Montiel & Mackenzie 2011).

**Figure 2.**
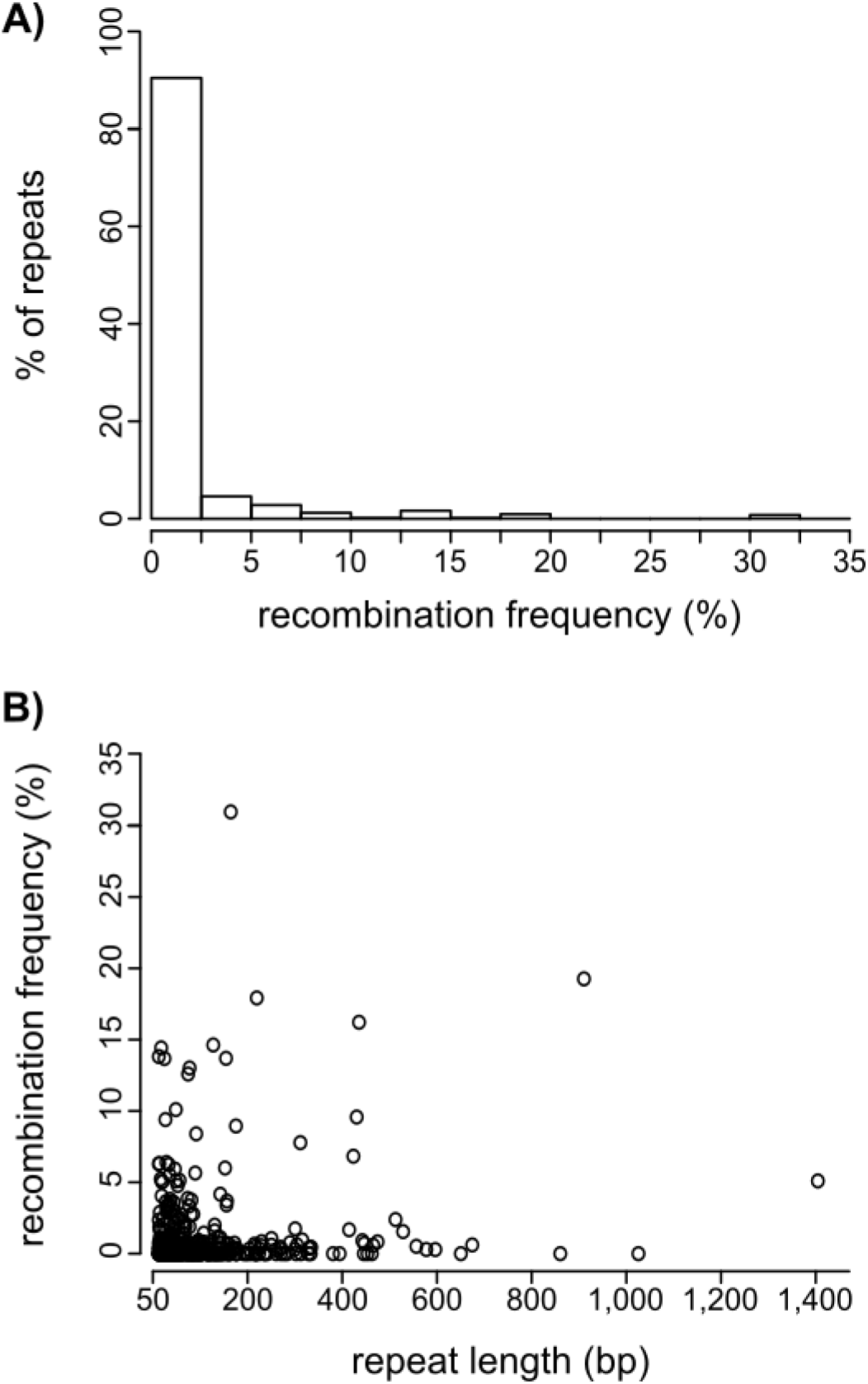
Recombination frequency at inverted repeats ≥ 50 bp with ≥ 80% pairwise identity as inferred from long reads mapping to expected recombination products (alternative genome configurations; AGCs). A) Most repeat pairs have little or no evidence of recombination, but a minority are highly active. B) Repeat length explains very little of the variation in recombination frequency (r^2^ = 0.03).

### A reappraisal of intramolecular mitogenome recombination

Plant mitogenomes are widely cited to undergo frequent, reciprocal recombination at large repeats and only rarely elsewhere based on the results of seminal pre-genomic studies (Lonsdale et al. 1984; Palmer & Shields 1984; Arrieta-Montiel & Mackenzie 2011). As our results suggested *P. abies* deviates from this pattern, we tested if this canonical model of mitogenome evolution is still supported by modern sequencing data. We searched the *ca*. 200 published tracheophyte mitogenomes and identified those that 1) analyzed and reported repeat-by-repeat recombination dynamics and 2) achieved at least a 60x average read depth of coverage. This coverage allows identification of AGCs comprising ≥ 1.6% of the read pool and establishes a baseline for comparing genomes sequenced to varying depths (e.g. 60 to over 1,000x), although confirmed AGCs have been documented at much lower frequencies (Woloszynska 2009; Arrieta-Montiel & Mackenzie 2011). Only 18 mitogenomes, representing ferns, gymnosperms, and four major angiosperm lineages, met both criteria.

Recombination patterns across tracheophytes provided mixed support for the canonical model of mitogenome evolution (Table 4, Table S5). Results are summarized as proportions in Table 4 to account for differences in repeat counts across species but are presented in absolute terms in Table S5. Consistent with the canonical model, large repeats tended to be more active than their smaller counterparts (Table 4). However, AGC abundances were not at equilibrium with their reciprocal configurations in most species (Table 4), indicating a lower recombination rate than anticipated from Southern blots (Maréchal & Brisson 2010). Although small repeats were overall less recombinogenic, active small repeats produced AGCs at similar frequencies as their larger counterparts in 33% of species, including *P. abies* (Table 4). All species with active small repeats except for *Silene conica* showed high AGC frequencies at repeats measuring just *ca*. 200 bp in length (Sloan et al. 2012; Mower, Floro, et al. 2012; Naito et al. 2013; Skippington et al. 2015; Pinard et al. 2019). While references to recombination rates are often necessarily vague, the 10-50% AGC abundances in these species exceed any reasonable interpretation of phrases such as ‘highly substoichiometric’ that are frequently used to describe the activity of small repeats (Woloszynska 2009; Arrieta-Montiel & Mackenzie 2011). Together, these studies point to the importance of small repeats as drivers of mitogenome evolution, a long anticipated (André et al. 1992) but rarely quantified phenomenon.

**Table 4.**
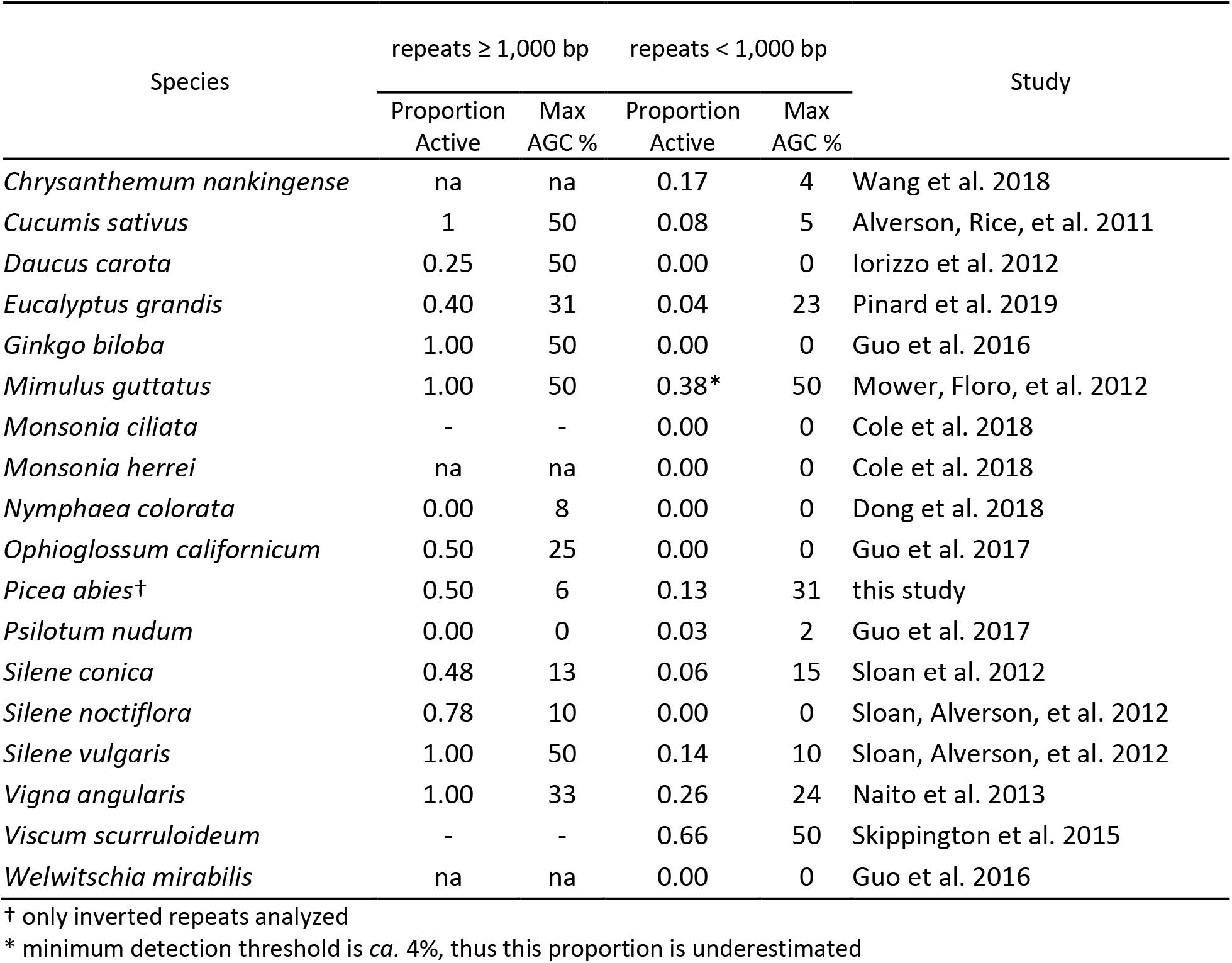
Recombination patterns summarized from 18 published vascular plant mitogenomes. ‘Proportion active’ refers to the fraction of repeats producing alternative genome configurations (AGCs) inferred to be the product of recombination in frequencies ≥ 1.6% of the parent molecule. ‘Max AGC’ denotes the maximum frequency obtained by any AGC in the given repeat size class. Missing data because repeats of a size class do not exist in a given genome are listed as ‘na’, whereas ‘-‘ indicates missing data due to study limitations.

Differences in nuclear-encoded repair pathways may be the mechanism underlying heterogeneity in recombination rates (Maréchal & Brisson 2010; Gualberto & Newton 2017). For example, *Arabidopsis* mutants for mitochondrial repair and surveillance genes recombined more readily at small repeats than their wild type counterparts (Zaegel et al. 2006; Shedge et al. 2007; Miller-Messmer et al. 2012; Wallet et al. 2015). While mutations associated with these or undiscovered genes could contribute to the diversity of recombination dynamics, the results in Table 4 do not show any discernable phylogenetic signal. If nuclear-encoded repair genes directly explain most of the observed differences in recombination dynamics, then this lack of signal implies repeated, independent evolution. Identifying factors influencing recombination rates, and in turn how – or if – their variation explains facets of genome evolution such as genome size, repeat content, and mutation rate would be aided by more consistent reporting of recombination rates at a minimum.

### Rampant mitogenome rearrangements in *Picea*

Rearrangements between the *P. abies* and *P. glauca* mitogenomes occurred an average of every 1,540 bp and blocks of synteny rarely extended beyond gene boundaries (Fig. 3). After accounting for the draft state of the genomes (Muñoz & Sankoff 2010), a parsimonious rearrangement scenario (Tesler 2002) required 1,292 events to explain the size and distribution of synteny blocks between the *Picea* species (1,310 events, unadjusted for assembly scaffold number). Assuming a divergence time of approximately 15 My ago (95% CI: 10-18 My; Feng et al. 2018) results in an absolute rearrangement rate of around 36 – 65 rearrangements My^−1^. This rate is similar to those observed in *Silene vulgaris* and some closely-related *Monsonia* species (Cole et al. 2018), which appear exceptionally rearranged relative to other eukaryotes. Rearrangement inference methods used here and by Cole et al. (2018) do not correct for events that have been lost due to the erosion of shared sequence over time, which should underestimate rearrangements given increasing divergence. When comparing similarly diverged mitogenomes (*ca*. 50% shared sequence) to mitigate this bias, the high rearrangement rate in *Picea* is even clearer: 43 *vs*. an average of 1 rearrangement My^−1^ (SD: 0.20) for four other species pairs (Table S6).

**Figure 3.**
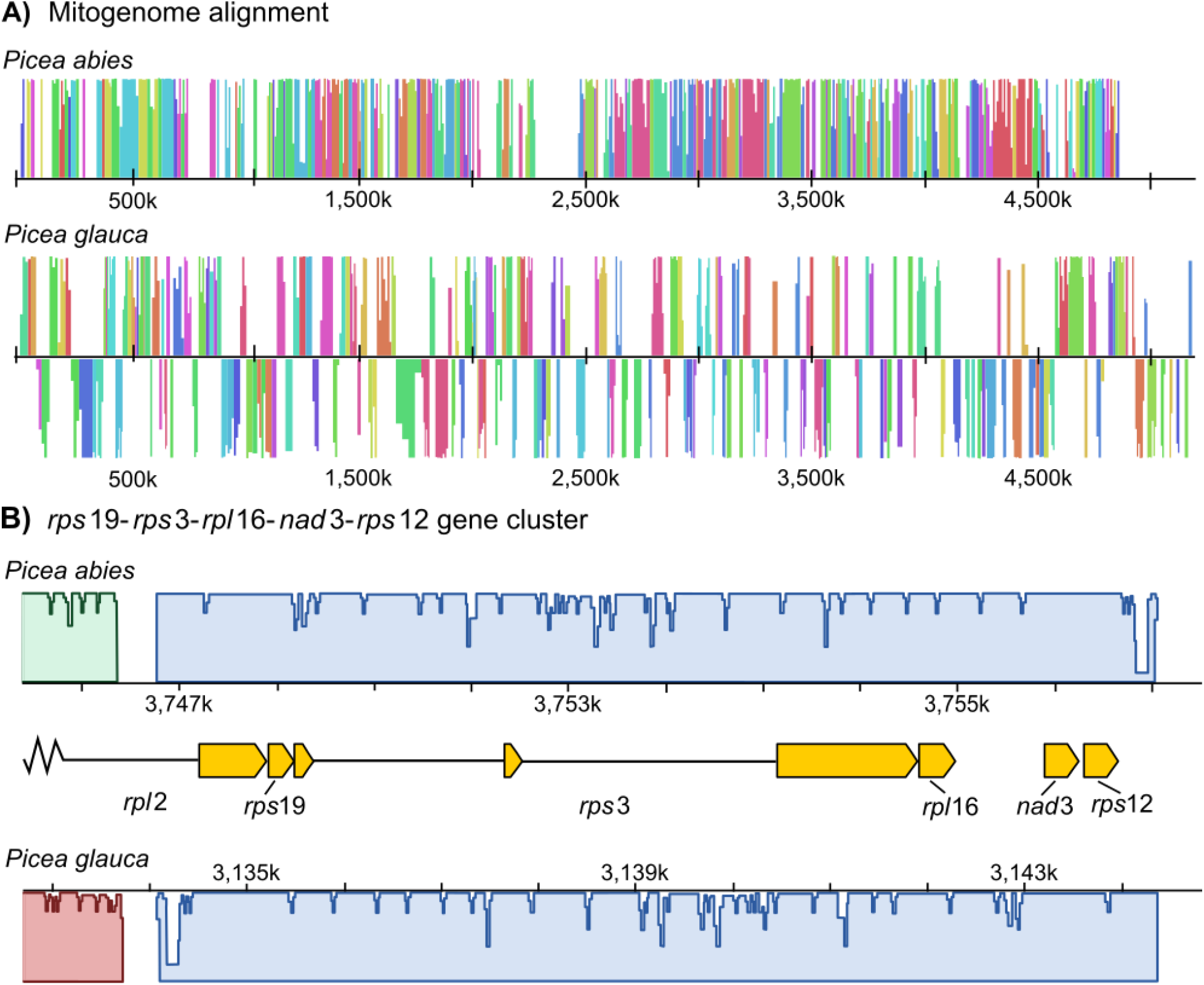
The mitogenomes of *Picea abies* and *P. glauca* are extensively rearranged and collinear regions are limited to genic regions. An absolute rearrangement rate of 36-65 My^−1^ is needed to explain this level of structural divergence. A) Simplified diagram of the *P. abies*-*P. glauca* mitogenome alignment, where colored blocks represent corresponding homologous regions free of internal re-arrangements (locally collinear blocks; LCBs) and heights are proportional to pairwise sequence identity. LCBs below the center line in *P. glauca* are inverted with respect to *P. abies*. White space indicates regions with no homology. Only LCBs longer than 2,000 bp are shown. B) Two gene clusters widely conserved in plant mitogenomes are also found in *Picea* within a 9 kb block, which also serves to illustrate the typical extent of synteny beyond genic regions. Gene structures are indicated by yellow boxes and introns by black lines.

Mitogenome shuffling in *Picea* contrasts with the remarkably static Pinaceae nuclear genome, where a high degree of synteny has persisted among genera after ≥ 140 My of divergence (Krutovsky et al. 2004; Pelgas et al. 2006; Pavy et al. 2012). Even the nuclear genomes of Cupressaceae and Pinaceae, which last shared a common ancestor in the Carboniferous (Leslie et al. 2018), are more collinear than the *P. abies* and *P. glauca* mitogenomes (Ehrenmann et al. 2015). Despite their extensive rearrangements, two of the three gene clusters widely preserved in tracheophytes from their α-proteobacterial endosymbiont (Dong et al. 2018) were also maintained in *Picea* within the same 9 kb block: *rpl*2-*rps*19-*rps*3-*rpl*16 and *nad*3-*rps*12 (Fig. 3B). The third widely-conserved gene cluster – *rrn*5-*rrn*18 – has not been preserved in *Picea*, *Pinus taeda*, or *Welwitschia* but persisted in *Cycas* and *Ginkgo* (Guo et al. 2016), suggesting it was lost in the common ancestor of conifers and gnetophytes.

Intramolecular recombination creates AGCs that can potentially be transmitted as rearrangements (Gualberto & Newton 2017; Cole et al. 2018). However, the path from isomer to rearrangement is not straightforward because mitogenomes are subject to genetic drift within an individual and within a population (Gualberto & Newton 2017). After an AGC is produced, it must survive multiple rounds of cell division, be recruited into the germline, and persist through potentially multiple generations before the rearrangement becomes firmly established within an individual (Davila et al. 2011). Proliferation throughout the population could then occur as other polymorphisms, probably predominately through stochastic demographic forces but possibly also through natural selection (Shedge et al. 2010). Several aspects of this process are unknown, including how mitogenomes are replicated (Gualberto & Newton 2017), the timing of germline segregation (Lanfear 2018), and when, where, and under what circumstances AGCs arise. For these reasons, the relationship between recombination within individuals summarized in Table 4 and re-arrangement rates among species is unclear. For example, *Monsonia* and *Silene vulgaris* have markedly different levels of intramolecular recombination (Table 4), yet have similar rearrangement rates (Cole et al. 2018).

### Sequence divergence in *Picea* and among gymnosperms

A strong dichotomy between rates of sequence and structural evolution is a well-known feature of plant mitogenomes (Palmer & Herbon 1988) and is also the dynamic found in *Picea* and other Pinaceae conifers (Fig. 4). Substitution rates across all protein-coding genes averaged 0.0023 for synonymous and 0.0012 for nonsynonymous sites. These rates are about 1/2 of those in the plastid genome (Sullivan et al. 2017) and 1/4 of the nuclear genome (Buschiazzo et al. 2012; Chen et al. 2012). Assuming a divergence time of 10 – 17 My between *P. abies* and *P. glauca* (Feng et al. 2018), mitochondrial substitution rates in *Picea* fall within the lower end of absolute rates observed in angiosperms (*dS* = 7.65 × 10^−11^; *dN* = 4.00 × 10^−11^ assuming a 15 My divergence, cf. Mower et al. 2007) and are slightly higher than in *Cycas* and *Ginkgo* (Guo et al. 2016). Absolute substitution rates in *Pinus*, however, include the lowest rates reported so far in vascular plants (Richardson et al. 2013). Using divergence times from Saladin et al. (2017), rates ranged from *d*S = 0.60 × 10^−11^ and *d*N = 0.30 × 10^−11^ site/year in *Pinus taeda* to *d*S = 3.15 × 10^−11^ and *d*N = 0.17 × 10^−11^ in *Pinus strobus*. Previous work with more taxa, but fewer loci, similarly inferred exceptionally low substitution rates in some *Pinus* species (Wang & Wang 2014). At the other extreme, absolute *d*S measured *ca*. 30.0 × 10^−11^ site/year in *Welwitschia* and *Araucaria heterophylla* when assuming a divergence from Pinaceae around 342 My ago (Lu et al. 2014). Relative substitution rates were consistent with previous studies analyzing fewer genes or species (e.g. Mower et al. 2007; Guo et al. 2016) and are reported in Table S7.

**Figure 4.**
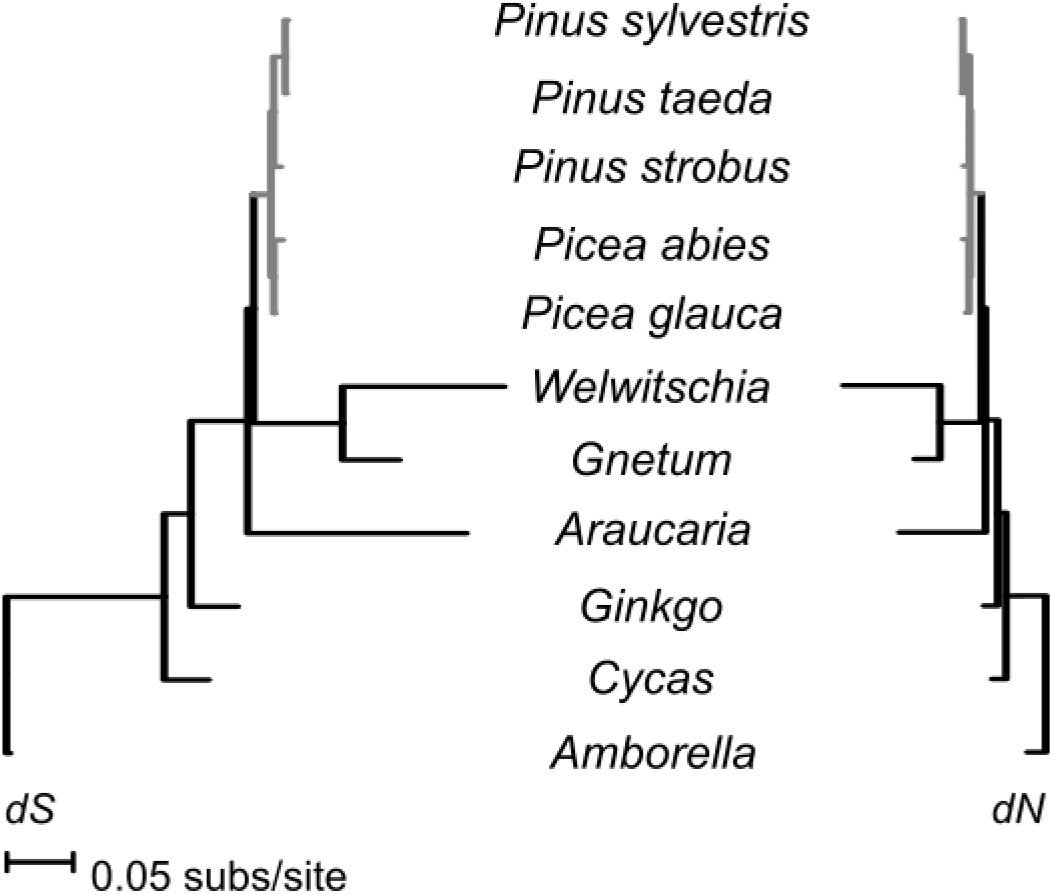
Mean substitution rates at synonymous (dS) and nonsynonymous (dN) sites vary 100-fold among gymnosperms. Rates within Pinaceae, in gray, are more consistent but are about 6-fold higher in *Picea* than in the 75% smaller *Pinus* mitogenomes.

### Intraspecific variation

We used the PacBio Sequel^®^ system to sequence partial mitogenomes from two *P. abies* on the alpine tundra in central Sweden, about 400 km southwest from the reference tree. *Picea* megafossil remains in this region date to *ca*. 11,000 ^14^C yr^−1^, which suggests the presence of high-latitude glacial refugia in Scandinavia (Kullman 2008). Both formed small clonal groups above the modern tree line and are spatially associated with megafossils dated to 5,120 and 4,820 ^14^C yr^−1^, respectively, raising the possibility that these clonal groups are extremely ancient (Kullman 2001). Thus, the mitogenomes of these two trees is relevant to post-glacial recolonization history and to the study of mitochondrial repair and recombination dynamics in long-lived organisms. In total, we recovered 499,180 and 553,660 bp, respectively, of which 82,443 was shared between the two trees. Nine variant sites were shared by the two trees relative to the reference tree. In contrast, 58 structural variations were found between these putatively ancient trees and the reference individual: 29 insertions, 17 deletions, 13 translocations, nine duplications, and seven inversions. More data is needed to address ecological and molecular hypotheses, but these sequences support a high rate of structural evolution in *Picea*.

## Conclusion

Plant mitogenomes are highly variable in size, repeat and gene content, and mutation and rearrangement rates. The underlying processes generating this heterogeneity are largely unclear: for each mitogenome supporting a given mechanistic hypothesis, another often suggests the opposite, such as in the relationship between mutation rate and genome size (cf. Sloan et al. 2012; Skippington et al. 2015, Christensen 2018). The *P. abies* mitogenome and our comparative analyses may lend support to the view of plant mitogenomes as highly idiosyncratic and driven mainly by rapid evolution and/or considerable genetic drift. As in previous studies, we found no clear relationship between genome size, repeat content, intergenomic transfer, or substitution rate. However, we identified recombination as an underinvestigated mechanism of plant mitogenome evolution, despite being recognized as a likely source of the differences among eukaryotes (Palmer & Herbon 1988; Gray et al. 1999). Recombination rates vary extensively among plants and small repeats (< 1,000 bp) are highly active in a third of the reported species. Recombination affects the accumulation of mutations (Maréchal & Brisson 2010; Christensen 2018), influences genome size (Christensen 2018), and induces structural rearrangements (Palmer & Herbon 1988), making this variation a potential but understudied contributor to the diversity of plant mitogenomes.

## Methods

### Machine learning classification of genomic scaffolds

The support vector machine involved four steps: 1) identification of high-quality training data, 2) training the classifier using this data and pre-determined genomic features, 3) evaluation of the model performance, and 4) applying the classifier to the *P. abies* v. 1.0 genome assembly. To curate a reliable set of positive and negative training data from *P. abies*, we first discarded scaffolds < 500 bp and masked repetitive elements using RepeatMasker v. open-4.0.1 (Smit et al. 2013). We aligned the remaining scaffolds to a set of relatively well-annotated genomes, including *Arabidopsis thaliana* and *Populus trichocarpa* and 18 additional organelle genomes (Table S8), using blastn and tblastx in BLAST version 2.2.29+ with default parameters except for an e-value cutoff of 1e-20. Known numts (nuclear mitochondrial DNA sequences) were masked from the *Arabidopsis* assemblies. *P. abies* sequences were considered robustly assigned if they matched an annotated gene region on the subject genome and 1) the top bitscore was > 100; 2) the quotient between the top bitscore and next-best hit was > 1.2; 3) alignments spanned at least 150 bp for blastn and 100 aa for tblastx; 4) blastn and tblastx annotations were identical if both existed; and 5) at least two different reference species contained the BLAST hit. Scaffolds meeting all these criteria were then classified as mitochondrial or non-mitochondrial and as training data for the SVM classifier. This yielded 2.0 Mb and 51.1 Mb of positive and negative data, respectively, for training the SVM.

After identifying robust positive and negative training datasets, we trained the SVM classifier to distinguish between them using pre-determined features. Following previous studies (Nystedt et al. 2013; Jackman et al. 2015), we used the standardized values of the natural logarithm of mean depth of coverage, the proportion of ambiguous nucleotides (%N), and the proportion of guanine and cytosine (%GC) for each scaffold, and a kmer-based score based on the frequency of nonredundant ‘mitochondrial-like’ or ‘non-mitochondrial-like’ oligonucleotide sequences in a scaffold. We calculated probability tables for occurring kmers based on positive and negative training sequences and calculated a score for each accordingly:

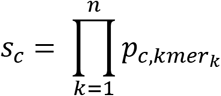

where *c* is the class, *n* is the number of kmers in the scaffold, and *p_c,kmer_k__* is the probability of kmer *k* occurring in a sequence of class *c*. The final kmer classifier score for each scaffold was calculated as:

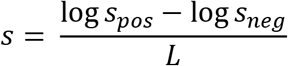

where *L* is the scaffold length and *s_pos_* and *s_neg_* are the positive and negative scores as defined above. Thus, smaller scores are consistent with plastid or nuclear sequence whereas larger scores indicate a scaffold comprising more mitochondrial-like kmers. Optimal kmer size for the classifier (k = 7, here) was assessed using test and training data from *A. thaliana* and visual inspection of the receiver operating characteristic (ROC) curves. After defining these features, we used SVMlight (Joachims 1999) with a Gaussian kernel to conduct the training step and to apply the learned model to the set of *P. abies* scaffolds with length > 500 bp and coverage > 100x.

We assessed the SVM model performance using cross-validation. The initial training data were divided randomly into nine trials varying in training subsets from 10-90%, each consisting of 100 cross-validation tests. Kmer scores, SVM training, and classification were carried out as described above. For each trial, we calculated the mean false discovery rate (FDR) and true positive recall.

### Mitogenome assembly

We screened subreads from 77 PacBio Sequel^®^ SMRT cells generated from genomic DNA (Street et al., unpublished data) for 27-mer matches to any of the SVM-classified scaffolds using BBDuk v. 35.14 (http://jgi.doe.gov/data-and-tools/bb-tools). Enriched reads were assembled using canu v. 1.7 (Koren et al. 2017), MECAT v. 1.3 (Xiao et al. 2017), and SMARTdenovo (https://github.com/ruanjue/smartdenovo). Canu was run using default parameters, except `corMaxEvidenceErate` was set to 0.15 as recommended in the manual for repetitive and GC-skewed genomes. MECAT was also run using default parameters using a target of 45x, 55x, and 65x coverage of the longest corrected reads. For MECAT and canu, we specified a genome size of approximately 5.0 Mb, which is used to estimate the coverage of the input reads. Because SMARTdenovo does not include a preassembly read correction step, we used the 45x, 55x, and 65x corrected reads from MECAT as input and the default parameters were then used for assembly.

We selected the most contiguous assembly from each assembler for further refinement. We passed these assemblies and their constituent reads to FinisherSC to reconstruct the overlap graph and identify contigs that can be robustly merged (Lam et al. 2015). Next, we used tblastx to identify the contig in each upgraded assembly containing the best match to each of the 41 protein-coding genes in the *Cycas* mitogenome (Chaw et al. 2008), which we retained as ‘high-confidence’ assemblies. Then, we used pairwise alignments from NUCmer (Kurtz et al. 2004) to break the three assemblies at major disagreements. The resulting contigs were put through the assembly reconciliation pipeline implemented in CISA v. 1.3 (Lin & Liao 2013) to re-assemble them into a single draft. After checking for circular contigs using dot plots, we evaluated the quality of the assembly by aligning the corrected reads using minimap2 (Li 2018) to the draft and visually inspected their congruency in IGV v. 2.4.14. (Thorvaldsdóttir et al. 2012). Finally, we used the partial order alignment graph approach implemented in Racon to re-call the consensus sequence from the minimap2 alignment (Vaser et al. 2017).

### Genome annotation

We reused repeat libraries curated for the *P. abies* v. 1.0 assembly (Nystedt et al. 2013) as input for RepeatMasker v. 4.0.7. (Smit et al. 2013) to identify TEs including long retrotransposons (LTRs), transposable elements, and other interspersed elements (LINEs and SINES). We additionally used RepeatModeler v. 1.0.8. to identify potential *de novo* repeats with significant similarity to the RepBase and Dfam databases (Smit et al. 2013). Direct and inverted repeats were identified with self *vs*. self blastn searches following Guo et al. (2016). Mitochondrial DNA of plastid origin (MIPTs) were identified following the methods of Guo et al. (2016) and shared mitochondrial-nuclear DNA following the methods of (Alverson, Rice, et al. 2011).

We used MAKER v. 3.01.2 to annotate protein-coding genes (Cantarel et al. 2008). Complex repeats identified by RepeatMasker and RepeatModeler were hard-masked prior to annotation, whereas simple repeats were re-annotated and soft masked by MAKER. As empirical gene evidence, we used the transcriptomes of *P. abies*, *P. glauca*, *P. sitchensis, Pinus pinaster*, *Pinus sylvestris*, and *Pinus taeda* from the PLAZA 3.0 database (Proost et al. 2014) and all curated plant sequences (Swiss-Prot) from UniProt as protein evidence. This provided a total of 312,953 ESTs and 37,954 proteins for use as evidence alignments. Although we initially attempted to use the SNAP and AUGUSTUS *ab initio* gene predictors, we found the number of high-confidence genes was insufficient for training. Therefore, annotations are based on empirical evidence alone. Coding sequences were manually curated to improve identification of reading-frames and gene structure based on sequence homology. Hypothetical proteins were compared to an RNA-Seq dataset obtained from needles (PRJEB26398, Schneider et al., in preparation), in which transcripts were extracted using a ribominus kit and subjected to *de novo* transcriptome assembly using Trinity v. 2.8.4 using default parameters (Grabherr et al. 2011). The *de novo* reconstructed transcripts were aligned to the *P. abies* genome v. 1.0 retrieved from ConGenIE.org (Sundell et al. 2015) using the GMAP v. 2015.11.20 (Wu & Watanabe 2005). The alignment coordinates were intersected with the ab-initio prediction to validate the predicted hypothetical proteins using an ad-hoc R script.

### Identification of repeat-mediated recombination and rearrangements

For each inverted repeat pair, we extracted the ± 2,000 bp single-copy flanking regions and constructed their expected recombination products for use as an alternative genome reference (Day & Madesis 2007, Fig. S2). We used minimap2 to map the enriched PacBio reads against these four sequences, with the expectation that reads spanning the repeat and flanking regions of the alternative configurations with robust mapping quality (MQ > 50) represent actual structural variations present within the sequenced individual tree (e.g. Park et al. 2014; Skippington et al. 2015; Cole et al. 2018; Dong et al. 2018).

We aligned the mitogenomes of *P. abies* and *P. glauca* using the rearrangement-aware progressiveMAUVE aligner (Darling et al. 2010), which identifies locally-collinear blocks (LCBs) internally free of recombination and reorders them against a selected genome. Then, we used GRIMM 2.01 (Tesler 2002) to infer a minimum, optimum rearrangement history, assuming signed undirected chromosomes. Apparent rearrangements introduced by the draft state of the genomes were corrected following Muñoz & Sankoff (2010) and Cole et al. (2018). Shared sequence between *P. abies* and *P. glauca* was estimated using blastn and the same parameters as Guo et al. (2016) for consistency. To put the re-arrangement rates inferred in *Picea* into a broader phylogenetic context, we re-aligned and estimated minimum numbers of rearrangements from the species pairs in Guo et al. (2016) with similar levels of divergence (Table S6).

We searched manuscripts associated with all 127 tracheophyte mitogenomes in the NCBI Genome resource (accessed March 11, 2019) for estimations of recombination rates. In addition, we exhaustively searched Google Scholar for ‘plant mitochondrial genomes’ for publications since 2011, the year of the last extensive review of plant mitogenome recombination (Alverson, Zhuo, et al. 2011). To be included in our analysis, we required 1) genomes to be assembled using high-throughput sequencing at a depth of coverage sufficient to allow detection of AGCs comprising ≥ 1.6% of the read pool, 2) estimates of repeat-by-repeat recombination rates inferred from mapping statistics or alignment rates to be clearly reported, either tabulated or in figures sufficiently detailed to allow precise interpolation of from vectorized figures using Inkscape v. 0.91 (e.g. Sloan et al. 2012, their Fig. S6). Wherever possible, we limited the analyses to repeats ≥ 50 bp with ≥ 80% identity, although some studies employed more stringent cut-offs. Relevant details for each genome, including the sequencing depth of coverage, repeat number, range of repeat sizes and identities, and other analytical details are summarized in Table S5.

### Comparative analysis of gymnosperm mitogenomes

We downloaded the mitogenomes of *Amborella trichopoda, Cycas taitungensis*, *Ginkgo biloba*, *Welwitschia mirabilis*, *P. glauca*, and *Pinus taeda* from Genbank to compare substitution rates, gene and repeat content, and genomic structure. For analyses of nucleotide substitution rates, we also used the previously-sequenced mitogenome coding sequences of *Gnetum gnemon*. *Pinus sylvestris*, and *Araucaria heterophylla* from NCBI Genbank. Given the draft state of the *P. glauca* mitogenome, we retained only contigs with conserved mitochondrial genes, which resulted in a 5.2 Mb assembly. We re-annotated repeats and intergenome sequence transfers as described for *P. abies* for consistency.

Nucleotide substitution rates were estimated from the 41 protein-coding genes in Pinaceae. Coding sequences were extracted, aligned with MUSCLE v. 3.8.42 (Edgar 2004), and manually verified to ensure correct reading frames and complete codons. RNA editing sites were predicted using the PREP-mt webserver (Mower 2005) using a cutoff value of 0.2 as in Guo et al. (2016). Edited nucleotide sequences were re-aligned using the translation alignment option in Geneious v. 11.1.4 and poorly aligned blocks of codons were removed using Gblocks v. 0.91b webserver (Castresana 2000). First and second codon positions were extracted using DnaSP v. 6 (Rozas et al. 2017) and the maximum likelihood phylogeny of the concatenated alignment was inferred using RAxML v. 8.2.12 (Stamatakis 2014) under the GTR-GAMMA model of nucleotide evolution. This phylogeny agreed with the prevailing consensus of gymnosperm evolution (Lu et al. 2014), so we estimated estimate branch lengths in units of synonymous (*d*S) and nonsynonymous (*d*N) substitution rates under the free-ratio branch model in PAML on this constrained topology (Yang 2007).

### Intraspecific mitogenome variation in *Picea abies*

We selected two putatively ancient trees from Kullman (2008) and collected newly-flushed buds. Buds were immediately placed in a solution comprising 10 mM MOPS, 5% (v/v) dimethylsulfoxide and 5% (w/v) glycerol at pH 7.3 and stored on dry ice and then at −80°C until isolation. In brief, intact mitochondria were isolated using centrifugation and extraneous DNA was degraded using DNase, and then mitochondrial DNA was isolated using a gram-negative bacteria genomic DNA purification kit (Supplementary Methods 1). DNA library construction and PacBio RS II sequencing was performed at DUKE Center for Genomic and Computational Biology according to manufacturer’s instructions.

PacBio sub-reads were checked for adapter contamination and then aligned to the whole *P. abies* v. 1.0 assembly containing the new *de novo* mitogenome using minimap2 (Li 2018). Scaffolds identified as mitochondrial-like by the SVM in the *P. abies* v. 1.0 assembly but were not contained within the *de novo* assembly were left in place, as they may represent numts. Bases were called using GATK v. 3.8.0 Haplotype Caller in gvcf mode followed by joint-genotyping in `includeNonVariantSites` mode (Van der Auwera et al. 2013). As GATK does not assign confidence metrics to non-variant sites, we considered those covered by 10 to 35 reads to be represented. Sites with higher coverage represent unresolved repeats. Structural variants were called with Sniffles v. 1.0.11 using default filtering settings (Sedlazeck et al. 2018). Each structural variant was then visually validated in IGV v. 2.4.14. (Thorvaldsdóttir et al. 2012).

## Acknowledgments

The authors acknowledge support from the National Genomics Infrastructure in Uppsala funded by Science for Life Laboratory, the Knut and Alice Wallenberg Foundation and the Swedish Research Council, and SNIC/Uppsala Multidisciplinary Center for Advanced Computational Science for assistance with massively parallel sequencing and access to the UPPMAX computational infrastructure, and the Danish Council for Independent Research – Natural Science (to IMM). NRS is supported by the Trees and Crops for the Future (TC4F) project. Three anonymous reviewers provided helpful comments that improved the clarity and accuracy of the manuscript.

