## Supplementary material for "The mitogenome of Norway spruce and a reappraisal of mitochondrial recombination in plants"

Table S1. Support vector machine (SVM) cross-validation results of mean recall and false discovery rate (FDR) estimated from 100 replicates using 10, 50, and 90% training data.

| | |
|---|---|
| Average recall rates | |
| 10% | 0.53 |
| 50% | 0.88 |
| 90% | 0.92 |
| Average FDRs | |
| 10% | 0.13 |
| 50% | 0.20 |
| 90% | 0.22 |

Table S2. Transcriptome support for hypothetical proteins predicted by MAKER. Transcripts were first assembled *de novo* and then mapped to the *P. abies* reference genome with GMAP.

| annotation | coverage | identity | matches | mismatches | indels |
|---|---|---|---|---|---|
| hypothetical protein 02 | 100 | 99.5 | 370 | 2 | 0 |
| hypothetical protein 03 | 99.8 | 99.5 | 1464 | 1 | 6 |
| hypothetical protein 05 | 100 | 99.6 | 275 | 0 | 1 |
| hypothetical protein 06 | 100 | 98.4 | 240 | 4 | 0 |
| hypothetical protein 07 | 100 | 99.5 | 792 | 0 | 4 |
| hypothetical protein 08 | 100 | 99.6 | 2924 | 1 | 11 |
| hypothetical protein 10 | 100 | 99.8 | 904 | 0 | 2 |
| hypothetical protein 11 | 100 | 99.8 | 4080 | 2 | 8 |
| hypothetical protein 12 | 99.9 | 99.2 | 2501 | 10 | 11 |
| hypothetical protein 13 | 100 | 100 | 210 | 0 | 0 |
| hypothetical protein 14 | 82.5 | 99.2 | 387 | 3 | 0 |
| hypothetical protein 16 | 97.6 | 99.7 | 3292 | 3 | 7 |
| hypothetical protein 17 | 100 | 99.7 | 3990 | 5 | 6 |
| hypothetical protein 19 | 42.2 | 97 | 97 | 3 | 0 |
| hypothetical protein 20 | 100 | 99.2 | 3406 | 18 | 8 |
| hypothetical protein 21 | 100 | 83 | 229 | 8 | 39 |
| hypothetical protein 22 | 100 | 98.4 | 2451 | 10 | 29 |
| hypothetical protein 24 | 100 | 99.8 | 402 | 1 | 0 |
| hypothetical protein 24 | 68.4 | 79.5 | 550 | 52 | 90 |
| hypothetical protein 25 | 100 | 99.7 | 787 | 2 | 0 |

Table S3. Mitogenome sizes by species and accession number, if available.

| Accession | Species | Genome size (bp) |
|---|---|---|
| NC_036945.1 | *Mielichhoferia elongata* | 100,342 |
| NC_024518.1 | *Buxbaumia aphylla* | 100,725 |

| | | |
|---|---|---|
| NC_031846.1 | *Zygodon viridissimus* | 103,098 |
| NC_024516.1 | *Callicladium imponens* | 103,830 |
| NC_024522.1 | *Orthotrichum stellatum* | 104,131 |
| NC_016121.1 | *Anomodon rugelii* | 104,239 |
| NC_021931.1 | *Anomodon attenuatus* | 104,252 |
| NC_034907.1 | *Stoneobryum mirum* | 104,345 |
| NC_031392.1 | *Stoneobryum bunyaense* | 104,352 |
| NC_031212.1 | *Brachythecium rivulare* | 104,460 |
| NC_024514.1 | *Ptychomnion cygnisetum* | 104,480 |
| NC_027974.1 | *Sanionia uncinata* | 104,497 |
| NC_031767.1 | *Nyholmiella obtusifolia* | 104,603 |
| NC_024517.1 | *Ulota hutchinsiae* | 104,608 |
| NC_029355.1 | *Orthotrichum macrocephalum* | 104,624 |
| NC_031393.1 | *Ulota crispa* | 104,636 |
| NC_031391.1 | *Nyholmiella gymnostoma* | 104,638 |
| NC_031394.1 | *Ulota phyllantha* | 104,671 |
| NC_031389.1 | *Orthotrichum bicolor* | 104,736 |
| NC_028191.1 | *Tetraplodon fuegianus* | 104,741 |
| NC_029356.1 | *Orthotrichum diaphanum* | 104,744 |
| NC_026121.1 | *Lewinskya speciosa* | 104,747 |
| NC_031390.1 | *Orthotrichum callistomum* | 104,785 |
| NC_028040.1 | *Chionoloma tenuirostre* | 105,001 |
| NC_024515.1 | *Climacium americanum* | 105,048 |
| NC_007945.1 | *Physcomitrella patens* | 105,340 |
| NC_024519.1 | *Bartramia pomiformis* | 106,198 |
| NC_027515.1 | *Syntrichia filaris* | 106,343 |
| NC_026891.1 | *Frisvollia varia* | 106,358 |
| NC_026540.1 | *Niphotrichum ericoides* | 106,727 |
| NC_026890.1 | *Niphotrichum elongatum* | 106,746 |
| NC_029452.1 | *Racomitrium lanuginosum* | 106,795 |
| NC_025931.1 | *Dilutineuron laevigatum* | 106,809 |
| NC_026784.1 | *Codriophorus aciculare* | 106,818 |
| NC_026975.1 | *Racomitrium emersum* | 107,186 |
| NC_026974.1 | *Bucklandiella orthotrichacea* | 107,215 |
| NC_024290.1 | *Tetraphis pellucida* | 107,730 |
| NC_024523.1 | *Funaria hygrometrica* | 109,586 |
| NC_024520.1 | *Atrichum angustatum* | 115,146 |
| NC_024521.1 | *Sphagnum palustre* | 141,276 |
| NC_037041.1 | *Tritomaria quinquedentata* | 142,510 |
| NC_016122.1 | *Treubia lacunosa* | 151,983 |
| NC_035978.1 | *Calypogeia arguta* | 159,061 |
| NC_035979.1 | *Calypogeia suecica* | 161,960 |
| NC_035980.1 | *Calypogeia neogaea* | 162,175 |
| NC_035977.1 | *Calypogeia integristipula* | 163,057 |

| | | |
|---|---|---|
| NC_026901.1 | *Aneura pinguis* | 165,603 |
| NC_013444.1 | *Pleurozia purpurea* | 168,526 |
| NC_012651.1 | *Nothoceros aenigmaticus* | 184,908 |
| NC_001660.1 | *Marchantia polymorpha* | 186,609 |
| NC_035345.1 | *Zostera marina* | 191,481 |
| NC_013765.1 | *Phaeoceros laevis* | 209,482 |
| NC_039757.1 | *Chrysanthemum boreale* | 211,002 |
| NC_016125.1 | *Brassica rapa* | 219,747 |
| NC_016123.1 | *Brassica juncea* | 219,766 |
| NC_008285.1 | *Brassica napus* | 221,853 |
| NC_017840.1 | *Spirodela polyrhiza* | 228,493 |
| NC_016120.1 | *Brassica carinata* | 232,241 |
| KP030753.1 | *Brassica nigra* | 232,407 |
| NC_031896.1 | *Sinapis arvensis* | 240,024 |
| NC_037476.1 | *Anthoceros angustus* | 242,410 |
| KF442616.1 | *Eruca vesicaria* | 247,696 |
| HM562727.1 | *Silene latifolia* | 253,413 |
| KT988071.2 | *Schrenkiella parvula* | 255,773 |
| NC_018551.1 | *Raphanus sativus* | 258,426 |
| AP017303.1 | *Ipomoea nil* | 265,768 |
| KT971339.1 | *Medicago truncatula* | 271,618 |
| KX063855.1 | *Diplostephium hartwegii* | 277,718 |
| JQ248574.1 | *Daucus carota* | 281,132 |
| NC_031164.1 | *Saccharum officinarum* | 300,784 |
| NC_023337.1 | *Helianthus annuus* | 300,945 |
| CM007908.1 | *Helianthus annuus* | 301,004 |
| CM009589.1 | *Rosa chinensis* | 313,448 |
| NC_041093.1 | *Chenopodium quinoa* | 315,003 |
| KU318712.1 | *Allium cepa* | 316,363 |
| NC_037070.1 | *Arabis alpina* | 323,159 |
| NC_035618.1 | *Spinacia oleracea* | 329,613 |
| KX641464.1 | *Lagerstroemia indica* | 333,948 |
| NC_035317.1 | *Stratiotes aloides* | 349,058 |
| KF709392.1 | *Ajuga reptans* | 352,069 |
| JF920286.1 | *Brassica oleracea* | 360,271 |
| NC_041177.1 | *Morus notabilis* | 362,069 |
| NC_015099.1 | *Beta vulgaris* | 364,950 |
| KU187967.1 | *Ziziphus jujuba* | 365,190 |
| BK010421 | *Arabidopsis thaliana* | 367,808 |
| NC_002511.2 | *Beta vulgaris* | 368,801 |
| GQ856147.1 | *Citrullus lanatus* | 379,236 |
| JN872551.2 | *Lotus japonicus* | 380,861 |
| NC_015994.1 | *Beta macrocarpa* | 385,220 |
| NC_036467.1 | *Nicotiana attenuata* | 394,341 |

| | | |
|---|---|---|
| NC_018554.1 | *Malus domestica* | 396,947 |
| | *Monsonia camdeboense* | 400,130 |
| NC_015121.1 | *Vigna radiata* | 401,262 |
| NC_039768.1 | *Glycine soja* | 402,545 |
| JX463295.1 | *Glycine max* | 402,558 |
| KJ820684.1 | *Batis maritima* | 403,930 |
| AP012599.1 | *Vigna angularis* | 404,466 |
| | *Monsonia ciliata* | 412,246 |
| KU310670.1 | *Cannabis sativa* | 415,602 |
| NC_035964.1 | *Solanum pennellii* | 423,596 |
| NC_016742.1 | *Millettia pinnata* | 425,718 |
| | *Silene vulgaris* | 427,000 |
| BA000042.1 | *Nicotiana tabacum* | 430,597 |
| NC_022714.1 | *Triticum timopheevii* | 443,419 |
| NC_035963.1 | *Solanum lycopersicum* | 446,257 |
| KC208619.1 | *Butomus umbellatus* | 450,826 |
| NC_036024.1 | *Triticum aestivum* | 452,526 |
| AP008982.1 | *Triticum aestivum* | 452,528 |
| NC_023338.1 | *Vaccinium macrocarpon* | 459,678 |
| NC_035962.1 | *Bupleurum falcatum* | 463,792 |
| NC_008360.1 | *Sorghum bicolor* | 468,628 |
| NC_039660.1 | *Ammopiptanthus mongolicus* | 475,396 |
| AP013107.1 | *Aegilops speltoides* | 476,091 |
| EU431224.1 | *Carica papaya* | 476,890 |
| NC_040010.1 | *Eucalyptus grandis* | 478,813 |
| NC_039596.1 | *Styphnolobium japonicum* | 484,916 |
| NC_011033.1 | *Oryza sativa* | 490,520 |
| NC_007886.1 | *Oryza sativa* | 491,515 |
| KT959112.1 | *Castilleja paramensis* | 495,499 |
| NC_023209.1 | *Salvia miltiorrhiza* | 499,236 |
| BA000024.1 | *Beta vulgaris* | 501,020 |
| KM207685.1 | *Hyoscyamus niger* | 501,401 |
| HQ874649.1 | *Ricinus communis* | 502,773 |
| NC_016741.1 | *Dorcoceras hygrometricum* | 510,519 |
| | *Monsonia vanderietiae* | 510,607 |
| KJ865410.1 | *Capsicum annuum* | 511,530 |
| NC_029816.1 | *Oryza minuta* | 515,022 |
| NC_040989.1 | *Eleusine indica* | 520,691 |
| KR610474.1 | *Nelumbo nucifera* | 524,797 |
| AP017300.1 | *Hordeum vulgare* | 525,599 |
| JN098455.1 | *Mimulus guttatus* | 525,671 |
| CP018169.1 | *Oryza sativa* | 527,116 |
| | *Zea luxurians* | 539,000 |
| KR559021.1 | *Heuchera parviflora* | 542,954 |

| | | |
|---|---|---|
| KJ485850.1 | *Rhazya stricta* | 548,608 |
| NC_021152.1 | *Liriodendron tulipifera* | 553,721 |
| NC_013816.1 | *Oryza rufipogon* | 559,045 |
| NC_029039.1 | *Viscum album* | 565,432 |
| NC_038053.1 | *Senna tora* | 566,589 |
| AY506529.1 | *Zea mays* | 569,630 |
| NC_035549.1 | *Hibiscus cannabinus* | 569,915 |
| | *Zea perennis* | 570,000 |
| | *Monsonia marlothii* | 571,776 |
| AFYB00000000.1 | *Solanum lycopersicum* | 575,198 |
| LT855379.1 | *Betula pendula* | 581,505 |
| | *Monsonia emarginata* | 581,703 |
| KC189947.1 | *Vicia faba* | 588,000 |
| NC_038052.1 | *Bombax ceiba* | 594,390 |
| NC_029693.1 | *Salix purpurea* | 598,970 |
| | *Monsonia herrei* | 604,880 |
| | *Monsonia speciosa* | 605,487 |
| NC_037468.1 | *Nymphaea colorata* | 617,195 |
| NC_037463.1 | *Citrus sinensis* | 640,906 |
| NC_035075.1 | *Gossypium davidsonii* | 644,311 |
| NC_035074.1 | *Gossypium thurberi* | 644,395 |
| KU056812.1 | *Salix suchowensis* | 644,437 |
| NC_035076.1 | *Gossypium trilobum* | 644,460 |
| NC_031323.1 | *Hesperelaea palmeri* | 658,522 |
| NC_027407.1 | *Gossypium harknessii* | 666,081 |
| NC_027406.1 | *Gossypium hirsutum* | 668,584 |
| NC_029998.1 | *Gossypium raimondii* | 676,078 |
| NC_028254.1 | *Gossypium barbadense* | 677,434 |
| JX999996.1 | *Lolium perenne* | 678,580 |
| NC_031696.1 | *Cocos nucifera* | 678,653 |
| NC_008332.1 | *Zea mays* | 680,603 |
| NC_022796.1 | *Asclepias syriaca* | 682,498 |
| NC_035073.1 | *Gossypium arboreum* | 687,482 |
| NC_040998.1 | *Acacia ligulata* | 698,138 |
| NC_008362.1 | *Tripsacum dactyloides* | 704,100 |
| JN375330.1 | *Phoenix dactylifera* | 715,001 |
| NC_039738.1 | *Leucaena trichandra* | 722,009 |
| NC_027000.1 | *Geranium maderense* | 737,091 |
| FM179380.1 | *Vitis vinifera complete* | 773,279 |
| NC_035157.1 | *Populus davidiana* | 779,361 |
| KT337313.1 | *Populus tremula* | 783,442 |
| NC_028329.1 | *Populus tremula* x *Populus alba* | 783,513 |
| NC_041085.1 | *Populus alba* | 838,420 |
| NC_034982.1 | *Utricularia reniformis* | 857,234 |

| | | |
|---|---|---|
| NC_014050.1 | *Cucurbita pepo* | 982,833 |
| NC_035958.1 | *Platycodon grandiflorus* | 1,249,590 |
| NC_016005.1 | *Cucumis sativus* | 1,555,940 |
| NC_031360.1 | *Corchorus olitorius* | 1,829,340 |
| NC_031359.1 | *Corchorus capsularis* | 1,999,600 |
| KT894204.1 | *Corchorus capsularis* | 1,999,602 |
| | *Cucumis melo* | 2,900,000 |
| | *Silene noctiflora* | 6,728,000 |
| | *Silene conica* | 11,318,000 |
| | **average genome size** | **520,581** |

Table S4. Alternative genome configurations (AGC) by repeat length. Read count indicates the total number of reads aligned to the four possible repeat configurations; product1 and product2 denote the two repeats ± 2,000 bp as found in the reference genome and presumably represent the most abundant configurations; ac1 and ac2 are the two reciprocal products produced by the recombination of the reference-state repeats; and ac prop (%) gives their combined proportion of the total read count.

| | | **read counts** | | | | |
|---|---|---|---|---|---|---|
| **length** | **total** | **product1** | **product2** | **ac1** | **ac2** | **ac prop (%)** |
| 50 | 267 | 144 | 123 | 0 | 0 | 0.00 |
| 50 | 243 | 123 | 120 | 0 | 0 | 0.00 |
| 50 | 352 | 216 | 136 | 0 | 0 | 0.00 |
| 50 | 292 | 129 | 163 | 0 | 0 | 0.00 |
| 50 | 270 | 144 | 126 | 0 | 0 | 0.00 |
| 50 | 215 | 125 | 90 | 0 | 0 | 0.00 |
| 50 | 237 | 113 | 124 | 0 | 0 | 0.00 |
| 50 | 235 | 104 | 131 | 0 | 0 | 0.00 |
| 51 | 301 | 144 | 157 | 0 | 0 | 0.00 |
| 51 | 256 | 137 | 119 | 0 | 0 | 0.00 |
| 51 | 271 | 124 | 147 | 0 | 0 | 0.00 |
| 51 | 279 | 182 | 97 | 0 | 0 | 0.00 |
| 51 | 307 | 123 | 184 | 0 | 0 | 0.00 |
| 51 | 355 | 181 | 174 | 0 | 0 | 0.00 |
| 51 | 292 | 155 | 137 | 0 | 0 | 0.00 |
| 52 | 234 | 130 | 104 | 0 | 0 | 0.00 |
| 52 | 264 | 148 | 116 | 0 | 0 | 0.00 |
| 52 | 323 | 179 | 144 | 0 | 0 | 0.00 |
| 52 | 337 | 136 | 201 | 0 | 0 | 0.00 |
| 53 | 222 | 113 | 109 | 0 | 0 | 0.00 |
| 53 | 200 | 98 | 102 | 0 | 0 | 0.00 |
| 54 | 291 | 120 | 171 | 0 | 0 | 0.00 |
| 54 | 245 | 134 | 111 | 0 | 0 | 0.00 |

| | | | | | | |
|---|---|---|---|---|---|---|
| 54 | 246 | 111 | 135 | 0 | 0 | 0.00 |
| 54 | 296 | 135 | 161 | 0 | 0 | 0.00 |
| 55 | 340 | 159 | 181 | 0 | 0 | 0.00 |
| 55 | 214 | 84 | 130 | 0 | 0 | 0.00 |
| 55 | 325 | 148 | 177 | 0 | 0 | 0.00 |
| 55 | 385 | 216 | 169 | 0 | 0 | 0.00 |
| 55 | 288 | 147 | 141 | 0 | 0 | 0.00 |
| 55 | 242 | 95 | 147 | 0 | 0 | 0.00 |
| 56 | 275 | 134 | 141 | 0 | 0 | 0.00 |
| 56 | 270 | 174 | 96 | 0 | 0 | 0.00 |
| 56 | 237 | 122 | 115 | 0 | 0 | 0.00 |
| 56 | 253 | 130 | 123 | 0 | 0 | 0.00 |
| 56 | 324 | 133 | 191 | 0 | 0 | 0.00 |
| 56 | 258 | 124 | 134 | 0 | 0 | 0.00 |
| 57 | 295 | 150 | 145 | 0 | 0 | 0.00 |
| 57 | 268 | 153 | 115 | 0 | 0 | 0.00 |
| 57 | 369 | 227 | 142 | 0 | 0 | 0.00 |
| 57 | 250 | 109 | 141 | 0 | 0 | 0.00 |
| 57 | 335 | 131 | 204 | 0 | 0 | 0.00 |
| 57 | 298 | 144 | 154 | 0 | 0 | 0.00 |
| 57 | 272 | 96 | 176 | 0 | 0 | 0.00 |
| 57 | 250 | 141 | 109 | 0 | 0 | 0.00 |
| 57 | 283 | 140 | 143 | 0 | 0 | 0.00 |
| 57 | 393 | 226 | 167 | 0 | 0 | 0.00 |
| 57 | 261 | 144 | 117 | 0 | 0 | 0.00 |
| 58 | 242 | 129 | 113 | 0 | 0 | 0.00 |
| 58 | 262 | 127 | 135 | 0 | 0 | 0.00 |
| 58 | 329 | 137 | 192 | 0 | 0 | 0.00 |
| 58 | 294 | 150 | 144 | 0 | 0 | 0.00 |
| 58 | 189 | 90 | 99 | 0 | 0 | 0.00 |
| 58 | 248 | 140 | 108 | 0 | 0 | 0.00 |
| 59 | 352 | 193 | 159 | 0 | 0 | 0.00 |
| 59 | 255 | 96 | 159 | 0 | 0 | 0.00 |
| 59 | 307 | 147 | 160 | 0 | 0 | 0.00 |
| 59 | 266 | 105 | 161 | 0 | 0 | 0.00 |
| 59 | 289 | 153 | 136 | 0 | 0 | 0.00 |
| 59 | 269 | 114 | 155 | 0 | 0 | 0.00 |
| 60 | 364 | 188 | 176 | 0 | 0 | 0.00 |
| 60 | 255 | 148 | 107 | 0 | 0 | 0.00 |
| 60 | 299 | 153 | 146 | 0 | 0 | 0.00 |
| 60 | 366 | 217 | 149 | 0 | 0 | 0.00 |
| 60 | 292 | 172 | 120 | 0 | 0 | 0.00 |
| 61 | 327 | 196 | 131 | 0 | 0 | 0.00 |
| 61 | 338 | 169 | 169 | 0 | 0 | 0.00 |

| | | | | | | |
|---|---|---|---|---|---|---|
| 62 | 290 | 158 | 132 | 0 | 0 | 0.00 |
| 62 | 321 | 178 | 143 | 0 | 0 | 0.00 |
| 62 | 236 | 112 | 124 | 0 | 0 | 0.00 |
| 63 | 277 | 152 | 125 | 0 | 0 | 0.00 |
| 63 | 282 | 155 | 127 | 0 | 0 | 0.00 |
| 63 | 241 | 126 | 115 | 0 | 0 | 0.00 |
| 63 | 286 | 129 | 157 | 0 | 0 | 0.00 |
| 63 | 251 | 113 | 138 | 0 | 0 | 0.00 |
| 63 | 278 | 126 | 152 | 0 | 0 | 0.00 |
| 64 | 207 | 101 | 106 | 0 | 0 | 0.00 |
| 64 | 371 | 176 | 195 | 0 | 0 | 0.00 |
| 64 | 264 | 159 | 105 | 0 | 0 | 0.00 |
| 64 | 298 | 176 | 122 | 0 | 0 | 0.00 |
| 64 | 253 | 140 | 113 | 0 | 0 | 0.00 |
| 64 | 223 | 104 | 119 | 0 | 0 | 0.00 |
| 64 | 280 | 149 | 131 | 0 | 0 | 0.00 |
| 64 | 312 | 136 | 176 | 0 | 0 | 0.00 |
| 64 | 276 | 134 | 142 | 0 | 0 | 0.00 |
| 65 | 254 | 121 | 133 | 0 | 0 | 0.00 |
| 65 | 234 | 115 | 119 | 0 | 0 | 0.00 |
| 66 | 383 | 185 | 198 | 0 | 0 | 0.00 |
| 66 | 288 | 176 | 112 | 0 | 0 | 0.00 |
| 66 | 228 | 121 | 107 | 0 | 0 | 0.00 |
| 66 | 250 | 117 | 133 | 0 | 0 | 0.00 |
| 66 | 263 | 142 | 121 | 0 | 0 | 0.00 |
| 66 | 270 | 142 | 128 | 0 | 0 | 0.00 |
| 66 | 348 | 179 | 169 | 0 | 0 | 0.00 |
| 66 | 323 | 174 | 149 | 0 | 0 | 0.00 |
| 66 | 254 | 157 | 97 | 0 | 0 | 0.00 |
| 66 | 265 | 147 | 118 | 0 | 0 | 0.00 |
| 67 | 246 | 109 | 137 | 0 | 0 | 0.00 |
| 67 | 274 | 126 | 148 | 0 | 0 | 0.00 |
| 67 | 256 | 108 | 148 | 0 | 0 | 0.00 |
| 67 | 250 | 127 | 123 | 0 | 0 | 0.00 |
| 68 | 214 | 127 | 87 | 0 | 0 | 0.00 |
| 68 | 314 | 132 | 182 | 0 | 0 | 0.00 |
| 68 | 360 | 124 | 236 | 0 | 0 | 0.00 |
| 68 | 325 | 119 | 206 | 0 | 0 | 0.00 |
| 68 | 237 | 109 | 128 | 0 | 0 | 0.00 |
| 69 | 286 | 173 | 113 | 0 | 0 | 0.00 |
| 69 | 247 | 138 | 109 | 0 | 0 | 0.00 |
| 70 | 438 | 235 | 203 | 0 | 0 | 0.00 |
| 70 | 244 | 116 | 128 | 0 | 0 | 0.00 |
| 70 | 288 | 141 | 147 | 0 | 0 | 0.00 |

| | | | | | | |
|---|---|---|---|---|---|---|
| 70 | 263 | 160 | 103 | 0 | 0 | 0.00 |
| 72 | 272 | 136 | 136 | 0 | 0 | 0.00 |
| 72 | 272 | 103 | 169 | 0 | 0 | 0.00 |
| 73 | 281 | 145 | 136 | 0 | 0 | 0.00 |
| 73 | 316 | 136 | 180 | 0 | 0 | 0.00 |
| 73 | 227 | 91 | 136 | 0 | 0 | 0.00 |
| 73 | 337 | 228 | 109 | 0 | 0 | 0.00 |
| 73 | 242 | 126 | 116 | 0 | 0 | 0.00 |
| 73 | 359 | 116 | 243 | 0 | 0 | 0.00 |
| 74 | 318 | 161 | 157 | 0 | 0 | 0.00 |
| 74 | 264 | 153 | 111 | 0 | 0 | 0.00 |
| 75 | 259 | 136 | 123 | 0 | 0 | 0.00 |
| 75 | 370 | 153 | 217 | 0 | 0 | 0.00 |
| 75 | 300 | 135 | 165 | 0 | 0 | 0.00 |
| 75 | 228 | 137 | 91 | 0 | 0 | 0.00 |
| 75 | 271 | 126 | 145 | 0 | 0 | 0.00 |
| 76 | 289 | 171 | 118 | 0 | 0 | 0.00 |
| 76 | 304 | 173 | 131 | 0 | 0 | 0.00 |
| 76 | 333 | 180 | 153 | 0 | 0 | 0.00 |
| 76 | 302 | 162 | 140 | 0 | 0 | 0.00 |
| 76 | 231 | 108 | 123 | 0 | 0 | 0.00 |
| 76 | 282 | 127 | 155 | 0 | 0 | 0.00 |
| 77 | 236 | 112 | 124 | 0 | 0 | 0.00 |
| 77 | 249 | 100 | 149 | 0 | 0 | 0.00 |
| 77 | 257 | 139 | 118 | 0 | 0 | 0.00 |
| 78 | 285 | 129 | 156 | 0 | 0 | 0.00 |
| 78 | 288 | 115 | 173 | 0 | 0 | 0.00 |
| 78 | 249 | 137 | 112 | 0 | 0 | 0.00 |
| 78 | 313 | 181 | 132 | 0 | 0 | 0.00 |
| 78 | 394 | 245 | 149 | 0 | 0 | 0.00 |
| 80 | 271 | 123 | 148 | 0 | 0 | 0.00 |
| 81 | 259 | 131 | 128 | 0 | 0 | 0.00 |
| 81 | 276 | 122 | 154 | 0 | 0 | 0.00 |
| 81 | 288 | 112 | 176 | 0 | 0 | 0.00 |
| 81 | 330 | 155 | 175 | 0 | 0 | 0.00 |
| 81 | 348 | 142 | 206 | 0 | 0 | 0.00 |
| 82 | 318 | 191 | 127 | 0 | 0 | 0.00 |
| 83 | 272 | 137 | 135 | 0 | 0 | 0.00 |
| 83 | 227 | 106 | 121 | 0 | 0 | 0.00 |
| 83 | 254 | 142 | 112 | 0 | 0 | 0.00 |
| 83 | 245 | 117 | 128 | 0 | 0 | 0.00 |
| 84 | 338 | 215 | 123 | 0 | 0 | 0.00 |
| 84 | 305 | 147 | 158 | 0 | 0 | 0.00 |
| 84 | 295 | 172 | 123 | 0 | 0 | 0.00 |

| | | | | | | |
|---|---|---|---|---|---|---|
| 84 | 273 | 109 | 164 | 0 | 0 | 0.00 |
| 85 | 266 | 118 | 148 | 0 | 0 | 0.00 |
| 85 | 206 | 98 | 108 | 0 | 0 | 0.00 |
| 86 | 359 | 184 | 175 | 0 | 0 | 0.00 |
| 87 | 321 | 110 | 211 | 0 | 0 | 0.00 |
| 88 | 315 | 172 | 143 | 0 | 0 | 0.00 |
| 88 | 270 | 98 | 172 | 0 | 0 | 0.00 |
| 89 | 213 | 110 | 103 | 0 | 0 | 0.00 |
| 90 | 297 | 128 | 169 | 0 | 0 | 0.00 |
| 90 | 241 | 141 | 100 | 0 | 0 | 0.00 |
| 91 | 255 | 116 | 139 | 0 | 0 | 0.00 |
| 91 | 287 | 136 | 151 | 0 | 0 | 0.00 |
| 92 | 244 | 101 | 143 | 0 | 0 | 0.00 |
| 92 | 279 | 149 | 130 | 0 | 0 | 0.00 |
| 92 | 228 | 114 | 114 | 0 | 0 | 0.00 |
| 93 | 297 | 121 | 176 | 0 | 0 | 0.00 |
| 93 | 256 | 114 | 142 | 0 | 0 | 0.00 |
| 93 | 265 | 148 | 117 | 0 | 0 | 0.00 |
| 94 | 291 | 151 | 140 | 0 | 0 | 0.00 |
| 94 | 301 | 147 | 154 | 0 | 0 | 0.00 |
| 96 | 243 | 129 | 114 | 0 | 0 | 0.00 |
| 97 | 252 | 120 | 132 | 0 | 0 | 0.00 |
| 98 | 366 | 193 | 173 | 0 | 0 | 0.00 |
| 99 | 254 | 163 | 91 | 0 | 0 | 0.00 |
| 100 | 265 | 115 | 150 | 0 | 0 | 0.00 |
| 100 | 241 | 130 | 111 | 0 | 0 | 0.00 |
| 100 | 322 | 172 | 150 | 0 | 0 | 0.00 |
| 102 | 284 | 136 | 148 | 0 | 0 | 0.00 |
| 102 | 273 | 167 | 106 | 0 | 0 | 0.00 |
| 103 | 316 | 212 | 104 | 0 | 0 | 0.00 |
| 108 | 249 | 140 | 109 | 0 | 0 | 0.00 |
| 109 | 309 | 157 | 152 | 0 | 0 | 0.00 |
| 109 | 315 | 178 | 137 | 0 | 0 | 0.00 |
| 110 | 269 | 144 | 125 | 0 | 0 | 0.00 |
| 110 | 284 | 166 | 118 | 0 | 0 | 0.00 |
| 111 | 266 | 151 | 115 | 0 | 0 | 0.00 |
| 111 | 313 | 150 | 163 | 0 | 0 | 0.00 |
| 112 | 344 | 131 | 213 | 0 | 0 | 0.00 |
| 113 | 258 | 152 | 106 | 0 | 0 | 0.00 |
| 113 | 264 | 139 | 125 | 0 | 0 | 0.00 |
| 115 | 325 | 156 | 169 | 0 | 0 | 0.00 |
| 118 | 333 | 160 | 173 | 0 | 0 | 0.00 |
| 118 | 233 | 150 | 83 | 0 | 0 | 0.00 |
| 118 | 333 | 226 | 107 | 0 | 0 | 0.00 |

| | | | | | | |
|---|---|---|---|---|---|---|
| 119 | 260 | 134 | 126 | 0 | 0 | 0.00 |
| 119 | 271 | 142 | 129 | 0 | 0 | 0.00 |
| 119 | 223 | 77 | 146 | 0 | 0 | 0.00 |
| 120 | 346 | 217 | 129 | 0 | 0 | 0.00 |
| 120 | 265 | 125 | 140 | 0 | 0 | 0.00 |
| 120 | 290 | 102 | 188 | 0 | 0 | 0.00 |
| 120 | 336 | 152 | 184 | 0 | 0 | 0.00 |
| 121 | 276 | 108 | 168 | 0 | 0 | 0.00 |
| 122 | 189 | 88 | 101 | 0 | 0 | 0.00 |
| 122 | 281 | 170 | 111 | 0 | 0 | 0.00 |
| 123 | 299 | 114 | 185 | 0 | 0 | 0.00 |
| 123 | 198 | 97 | 101 | 0 | 0 | 0.00 |
| 123 | 198 | 103 | 95 | 0 | 0 | 0.00 |
| 124 | 350 | 130 | 220 | 0 | 0 | 0.00 |
| 126 | 348 | 165 | 183 | 0 | 0 | 0.00 |
| 128 | 215 | 132 | 83 | 0 | 0 | 0.00 |
| 128 | 322 | 148 | 174 | 0 | 0 | 0.00 |
| 129 | 241 | 131 | 110 | 0 | 0 | 0.00 |
| 130 | 228 | 123 | 105 | 0 | 0 | 0.00 |
| 130 | 235 | 140 | 95 | 0 | 0 | 0.00 |
| 131 | 254 | 115 | 139 | 0 | 0 | 0.00 |
| 132 | 320 | 159 | 161 | 0 | 0 | 0.00 |
| 133 | 274 | 135 | 139 | 0 | 0 | 0.00 |
| 134 | 235 | 116 | 119 | 0 | 0 | 0.00 |
| 134 | 297 | 180 | 117 | 0 | 0 | 0.00 |
| 134 | 210 | 98 | 112 | 0 | 0 | 0.00 |
| 136 | 219 | 112 | 107 | 0 | 0 | 0.00 |
| 137 | 254 | 110 | 144 | 0 | 0 | 0.00 |
| 137 | 242 | 105 | 137 | 0 | 0 | 0.00 |
| 138 | 281 | 139 | 142 | 0 | 0 | 0.00 |
| 139 | 276 | 141 | 135 | 0 | 0 | 0.00 |
| 140 | 306 | 99 | 207 | 0 | 0 | 0.00 |
| 141 | 197 | 105 | 92 | 0 | 0 | 0.00 |
| 141 | 195 | 91 | 104 | 0 | 0 | 0.00 |
| 151 | 293 | 141 | 152 | 0 | 0 | 0.00 |
| 152 | 260 | 107 | 153 | 0 | 0 | 0.00 |
| 152 | 192 | 102 | 90 | 0 | 0 | 0.00 |
| 154 | 284 | 131 | 153 | 0 | 0 | 0.00 |
| 155 | 237 | 127 | 110 | 0 | 0 | 0.00 |
| 155 | 279 | 150 | 129 | 0 | 0 | 0.00 |
| 157 | 226 | 96 | 130 | 0 | 0 | 0.00 |
| 157 | 289 | 158 | 131 | 0 | 0 | 0.00 |
| 157 | 250 | 132 | 118 | 0 | 0 | 0.00 |
| 161 | 251 | 120 | 131 | 0 | 0 | 0.00 |

| | | | | | | |
|---|---|---|---|---|---|---|
| 162 | 209 | 111 | 98 | 0 | 0 | 0.00 |
| 165 | 215 | 100 | 115 | 0 | 0 | 0.00 |
| 165 | 250 | 107 | 143 | 0 | 0 | 0.00 |
| 169 | 251 | 143 | 108 | 0 | 0 | 0.00 |
| 169 | 252 | 98 | 154 | 0 | 0 | 0.00 |
| 170 | 235 | 123 | 112 | 0 | 0 | 0.00 |
| 171 | 282 | 101 | 181 | 0 | 0 | 0.00 |
| 174 | 232 | 131 | 101 | 0 | 0 | 0.00 |
| 175 | 210 | 89 | 121 | 0 | 0 | 0.00 |
| 175 | 258 | 137 | 121 | 0 | 0 | 0.00 |
| 178 | 200 | 102 | 98 | 0 | 0 | 0.00 |
| 180 | 246 | 118 | 128 | 0 | 0 | 0.00 |
| 181 | 258 | 104 | 154 | 0 | 0 | 0.00 |
| 184 | 250 | 150 | 100 | 0 | 0 | 0.00 |
| 184 | 232 | 138 | 94 | 0 | 0 | 0.00 |
| 188 | 301 | 123 | 178 | 0 | 0 | 0.00 |
| 195 | 235 | 115 | 120 | 0 | 0 | 0.00 |
| 195 | 239 | 120 | 119 | 0 | 0 | 0.00 |
| 198 | 254 | 152 | 102 | 0 | 0 | 0.00 |
| 200 | 261 | 116 | 145 | 0 | 0 | 0.00 |
| 201 | 273 | 149 | 124 | 0 | 0 | 0.00 |
| 213 | 271 | 136 | 135 | 0 | 0 | 0.00 |
| 214 | 241 | 103 | 138 | 0 | 0 | 0.00 |
| 214 | 258 | 117 | 141 | 0 | 0 | 0.00 |
| 229 | 261 | 99 | 162 | 0 | 0 | 0.00 |
| 234 | 250 | 117 | 133 | 0 | 0 | 0.00 |
| 238 | 267 | 147 | 120 | 0 | 0 | 0.00 |
| 245 | 216 | 94 | 122 | 0 | 0 | 0.00 |
| 254 | 254 | 111 | 143 | 0 | 0 | 0.00 |
| 255 | 283 | 154 | 129 | 0 | 0 | 0.00 |
| 255 | 276 | 133 | 143 | 0 | 0 | 0.00 |
| 255 | 228 | 121 | 107 | 0 | 0 | 0.00 |
| 262 | 203 | 133 | 70 | 0 | 0 | 0.00 |
| 273 | 283 | 161 | 122 | 0 | 0 | 0.00 |
| 274 | 237 | 100 | 137 | 0 | 0 | 0.00 |
| 279 | 274 | 122 | 152 | 0 | 0 | 0.00 |
| 298 | 372 | 155 | 217 | 0 | 0 | 0.00 |
| 308 | 317 | 195 | 122 | 0 | 0 | 0.00 |
| 315 | 225 | 117 | 108 | 0 | 0 | 0.00 |
| 321 | 273 | 118 | 155 | 0 | 0 | 0.00 |
| 340 | 276 | 130 | 146 | 0 | 0 | 0.00 |
| 350 | 270 | 157 | 113 | 0 | 0 | 0.00 |
| 366 | 314 | 175 | 139 | 0 | 0 | 0.00 |
| 371 | 318 | 137 | 181 | 0 | 0 | 0.00 |

| | | | | | | |
|---|---|---|---|---|---|---|
| 418 | 255 | 102 | 153 | 0 | 0 | 0.00 |
| 432 | 342 | 203 | 139 | 0 | 0 | 0.00 |
| 483 | 291 | 143 | 148 | 0 | 0 | 0.00 |
| 491 | 232 | 112 | 120 | 0 | 0 | 0.00 |
| 500 | 203 | 112 | 91 | 0 | 0 | 0.00 |
| 688 | 217 | 74 | 143 | 0 | 0 | 0.00 |
| 898 | 307 | 120 | 187 | 0 | 0 | 0.00 |
| 1063 | 230 | 127 | 103 | 0 | 0 | 0.00 |
| 61 | 438 | 156 | 281 | 1 | 0 | 0.23 |
| 72 | 426 | 281 | 144 | 0 | 1 | 0.23 |
| 109 | 419 | 140 | 278 | 1 | 0 | 0.24 |
| 52 | 392 | 184 | 207 | 1 | 0 | 0.26 |
| 70 | 391 | 200 | 190 | 1 | 0 | 0.26 |
| 80 | 381 | 145 | 235 | 1 | 0 | 0.26 |
| 64 | 380 | 158 | 221 | 1 | 0 | 0.26 |
| 72 | 375 | 219 | 155 | 0 | 1 | 0.27 |
| 298 | 374 | 217 | 156 | 0 | 1 | 0.27 |
| 55 | 369 | 143 | 225 | 1 | 0 | 0.27 |
| 61 | 368 | 170 | 197 | 1 | 0 | 0.27 |
| 102 | 361 | 207 | 153 | 1 | 0 | 0.28 |
| 61 | 360 | 213 | 146 | 0 | 1 | 0.28 |
| 69 | 360 | 192 | 167 | 0 | 1 | 0.28 |
| 67 | 356 | 174 | 181 | 0 | 1 | 0.28 |
| 51 | 349 | 220 | 128 | 0 | 1 | 0.29 |
| 69 | 349 | 204 | 144 | 0 | 1 | 0.29 |
| 144 | 347 | 149 | 197 | 1 | 0 | 0.29 |
| 634 | 345 | 146 | 198 | 0 | 1 | 0.29 |
| 52 | 339 | 122 | 216 | 1 | 0 | 0.29 |
| 615 | 339 | 198 | 140 | 1 | 0 | 0.29 |
| 142 | 337 | 162 | 174 | 1 | 0 | 0.30 |
| 50 | 334 | 203 | 130 | 1 | 0 | 0.30 |
| 66 | 333 | 181 | 151 | 0 | 1 | 0.30 |
| 79 | 329 | 205 | 123 | 0 | 1 | 0.30 |
| 122 | 328 | 177 | 150 | 1 | 0 | 0.30 |
| 71 | 327 | 192 | 134 | 0 | 1 | 0.31 |
| 337 | 324 | 204 | 119 | 0 | 1 | 0.31 |
| 70 | 322 | 115 | 206 | 1 | 0 | 0.31 |
| 132 | 322 | 118 | 203 | 0 | 1 | 0.31 |
| 201 | 318 | 167 | 150 | 1 | 0 | 0.31 |
| 241 | 318 | 172 | 145 | 0 | 1 | 0.31 |
| 101 | 317 | 148 | 168 | 1 | 0 | 0.32 |
| 101 | 316 | 127 | 188 | 1 | 0 | 0.32 |
| 105 | 316 | 133 | 182 | 0 | 1 | 0.32 |
| 65 | 315 | 183 | 131 | 1 | 0 | 0.32 |

| | | | | | | |
|---|---|---|---|---|---|---|
| 51 | 314 | 157 | 156 | 1 | 0 | 0.32 |
| 51 | 312 | 183 | 128 | 0 | 1 | 0.32 |
| 53 | 308 | 128 | 179 | 1 | 0 | 0.32 |
| 132 | 308 | 156 | 151 | 1 | 0 | 0.32 |
| 307 | 308 | 121 | 186 | 0 | 1 | 0.32 |
| 53 | 306 | 180 | 125 | 1 | 0 | 0.33 |
| 56 | 306 | 166 | 139 | 1 | 0 | 0.33 |
| 137 | 305 | 144 | 160 | 1 | 0 | 0.33 |
| 56 | 302 | 138 | 163 | 1 | 0 | 0.33 |
| 92 | 302 | 148 | 153 | 1 | 0 | 0.33 |
| 118 | 302 | 186 | 115 | 1 | 0 | 0.33 |
| 94 | 301 | 170 | 130 | 1 | 0 | 0.33 |
| 65 | 300 | 192 | 107 | 0 | 1 | 0.33 |
| 84 | 300 | 126 | 173 | 1 | 0 | 0.33 |
| 122 | 300 | 151 | 148 | 0 | 1 | 0.33 |
| 108 | 299 | 158 | 140 | 0 | 1 | 0.33 |
| 59 | 298 | 107 | 190 | 1 | 0 | 0.34 |
| 110 | 298 | 145 | 152 | 1 | 0 | 0.34 |
| 58 | 297 | 146 | 150 | 0 | 1 | 0.34 |
| 60 | 293 | 153 | 139 | 0 | 1 | 0.34 |
| 111 | 293 | 200 | 92 | 0 | 1 | 0.34 |
| 54 | 292 | 191 | 100 | 1 | 0 | 0.34 |
| 54 | 292 | 167 | 124 | 1 | 0 | 0.34 |
| 130 | 291 | 113 | 177 | 1 | 0 | 0.34 |
| 148 | 290 | 130 | 159 | 1 | 0 | 0.34 |
| 75 | 288 | 127 | 160 | 1 | 0 | 0.35 |
| 78 | 288 | 110 | 177 | 0 | 1 | 0.35 |
| 57 | 287 | 185 | 101 | 1 | 0 | 0.35 |
| 120 | 287 | 186 | 100 | 1 | 0 | 0.35 |
| 81 | 286 | 155 | 130 | 0 | 1 | 0.35 |
| 116 | 286 | 115 | 170 | 1 | 0 | 0.35 |
| 131 | 286 | 155 | 130 | 1 | 0 | 0.35 |
| 152 | 285 | 155 | 129 | 0 | 1 | 0.35 |
| 75 | 284 | 138 | 145 | 0 | 1 | 0.35 |
| 71 | 283 | 111 | 171 | 1 | 0 | 0.35 |
| 262 | 283 | 129 | 153 | 1 | 0 | 0.35 |
| 111 | 282 | 130 | 151 | 1 | 0 | 0.35 |
| 141 | 282 | 175 | 106 | 0 | 1 | 0.35 |
| 117 | 280 | 158 | 121 | 1 | 0 | 0.36 |
| 206 | 280 | 153 | 126 | 1 | 0 | 0.36 |
| 87 | 279 | 139 | 139 | 0 | 1 | 0.36 |
| 106 | 278 | 149 | 128 | 0 | 1 | 0.36 |
| 213 | 278 | 131 | 146 | 1 | 0 | 0.36 |
| 57 | 277 | 115 | 161 | 1 | 0 | 0.36 |

| | | | | | | |
|---|---|---|---|---|---|---|
| 72 | 275 | 140 | 134 | 0 | 1 | 0.36 |
| 79 | 275 | 149 | 125 | 0 | 1 | 0.36 |
| 104 | 275 | 146 | 128 | 1 | 0 | 0.36 |
| 120 | 275 | 116 | 158 | 1 | 0 | 0.36 |
| 75 | 272 | 146 | 125 | 0 | 1 | 0.37 |
| 79 | 271 | 158 | 112 | 0 | 1 | 0.37 |
| 109 | 271 | 134 | 136 | 0 | 1 | 0.37 |
| 58 | 270 | 133 | 136 | 1 | 0 | 0.37 |
| 111 | 270 | 109 | 160 | 1 | 0 | 0.37 |
| 95 | 269 | 130 | 138 | 0 | 1 | 0.37 |
| 74 | 268 | 130 | 137 | 1 | 0 | 0.37 |
| 86 | 267 | 139 | 127 | 0 | 1 | 0.37 |
| 133 | 267 | 154 | 112 | 0 | 1 | 0.37 |
| 66 | 266 | 118 | 147 | 1 | 0 | 0.38 |
| 109 | 265 | 141 | 123 | 0 | 1 | 0.38 |
| 75 | 264 | 131 | 132 | 0 | 1 | 0.38 |
| 208 | 264 | 104 | 159 | 0 | 1 | 0.38 |
| 127 | 261 | 123 | 137 | 0 | 1 | 0.38 |
| 367 | 261 | 115 | 145 | 1 | 0 | 0.38 |
| 178 | 259 | 157 | 101 | 0 | 1 | 0.39 |
| 50 | 258 | 121 | 136 | 1 | 0 | 0.39 |
| 140 | 256 | 146 | 109 | 0 | 1 | 0.39 |
| 214 | 256 | 131 | 124 | 1 | 0 | 0.39 |
| 77 | 255 | 111 | 143 | 0 | 1 | 0.39 |
| 61 | 254 | 133 | 120 | 1 | 0 | 0.39 |
| 138 | 254 | 134 | 119 | 1 | 0 | 0.39 |
| 129 | 253 | 137 | 115 | 0 | 1 | 0.40 |
| 55 | 251 | 138 | 112 | 0 | 1 | 0.40 |
| 146 | 249 | 126 | 122 | 0 | 1 | 0.40 |
| 56 | 248 | 119 | 128 | 1 | 0 | 0.40 |
| 70 | 247 | 139 | 107 | 0 | 1 | 0.40 |
| 153 | 246 | 117 | 128 | 0 | 1 | 0.41 |
| 51 | 245 | 115 | 129 | 0 | 1 | 0.41 |
| 153 | 245 | 130 | 114 | 0 | 1 | 0.41 |
| 138 | 244 | 101 | 142 | 0 | 1 | 0.41 |
| 66 | 241 | 122 | 118 | 1 | 0 | 0.41 |
| 52 | 240 | 119 | 120 | 1 | 0 | 0.42 |
| 64 | 239 | 133 | 105 | 0 | 1 | 0.42 |
| 60 | 238 | 116 | 121 | 0 | 1 | 0.42 |
| 50 | 237 | 78 | 158 | 1 | 0 | 0.42 |
| 70 | 237 | 122 | 114 | 1 | 0 | 0.42 |
| 83 | 237 | 125 | 111 | 1 | 0 | 0.42 |
| 94 | 235 | 116 | 118 | 1 | 0 | 0.43 |
| 54 | 231 | 113 | 117 | 1 | 0 | 0.43 |

| | | | | | | |
|---|---|---|---|---|---|---|
| 71 | 230 | 137 | 92 | 0 | 1 | 0.43 |
| 92 | 227 | 113 | 113 | 0 | 1 | 0.44 |
| 84 | 226 | 101 | 124 | 0 | 1 | 0.44 |
| 372 | 225 | 107 | 117 | 0 | 1 | 0.44 |
| 84 | 224 | 122 | 101 | 1 | 0 | 0.45 |
| 89 | 224 | 92 | 131 | 1 | 0 | 0.45 |
| 64 | 223 | 119 | 103 | 0 | 1 | 0.45 |
| 370 | 223 | 120 | 102 | 0 | 1 | 0.45 |
| 101 | 221 | 141 | 79 | 1 | 0 | 0.45 |
| 116 | 221 | 104 | 116 | 1 | 0 | 0.45 |
| 63 | 220 | 109 | 110 | 0 | 1 | 0.45 |
| 133 | 220 | 107 | 112 | 0 | 1 | 0.45 |
| 307 | 219 | 102 | 116 | 1 | 0 | 0.46 |
| 132 | 218 | 110 | 107 | 1 | 0 | 0.46 |
| 253 | 211 | 108 | 102 | 0 | 1 | 0.47 |
| 92 | 209 | 99 | 109 | 1 | 0 | 0.48 |
| 51 | 391 | 181 | 208 | 2 | 0 | 0.51 |
| 594 | 192 | 101 | 90 | 1 | 0 | 0.52 |
| 54 | 362 | 186 | 174 | 0 | 2 | 0.55 |
| 107 | 358 | 133 | 223 | 1 | 1 | 0.56 |
| 266 | 353 | 162 | 189 | 1 | 1 | 0.57 |
| 69 | 352 | 148 | 202 | 1 | 1 | 0.57 |
| 109 | 347 | 216 | 129 | 0 | 2 | 0.58 |
| 61 | 346 | 156 | 188 | 2 | 0 | 0.58 |
| 53 | 345 | 188 | 155 | 2 | 0 | 0.58 |
| 83 | 340 | 179 | 159 | 0 | 2 | 0.59 |
| 77 | 337 | 189 | 146 | 2 | 0 | 0.59 |
| 503 | 168 | 67 | 100 | 0 | 1 | 0.60 |
| 288 | 334 | 155 | 177 | 1 | 1 | 0.60 |
| 342 | 323 | 119 | 202 | 0 | 2 | 0.62 |
| 712 | 322 | 135 | 185 | 1 | 1 | 0.62 |
| 69 | 318 | 111 | 205 | 0 | 2 | 0.63 |
| 79 | 313 | 170 | 141 | 1 | 1 | 0.64 |
| 90 | 311 | 136 | 173 | 0 | 2 | 0.64 |
| 198 | 304 | 135 | 167 | 1 | 1 | 0.66 |
| 76 | 299 | 148 | 149 | 2 | 0 | 0.67 |
| 79 | 299 | 154 | 143 | 2 | 0 | 0.67 |
| 153 | 298 | 122 | 174 | 0 | 2 | 0.67 |
| 252 | 297 | 165 | 130 | 0 | 2 | 0.67 |
| 58 | 296 | 145 | 149 | 2 | 0 | 0.68 |
| 79 | 295 | 180 | 113 | 1 | 1 | 0.68 |
| 62 | 294 | 132 | 160 | 2 | 0 | 0.68 |
| 82 | 293 | 130 | 161 | 2 | 0 | 0.68 |
| 109 | 293 | 167 | 124 | 1 | 1 | 0.68 |

| | | | | | | |
|---|---|---|---|---|---|---|
| 62 | 430 | 206 | 221 | 3 | 0 | 0.70 |
| 68 | 284 | 122 | 160 | 1 | 1 | 0.70 |
| 138 | 283 | 174 | 107 | 0 | 2 | 0.71 |
| 179 | 280 | 128 | 150 | 1 | 1 | 0.71 |
| 53 | 279 | 131 | 146 | 1 | 1 | 0.72 |
| 484 | 279 | 109 | 168 | 1 | 1 | 0.72 |
| 93 | 277 | 108 | 167 | 1 | 1 | 0.72 |
| 194 | 277 | 122 | 153 | 1 | 1 | 0.72 |
| 53 | 272 | 147 | 123 | 2 | 0 | 0.74 |
| 62 | 271 | 129 | 140 | 2 | 0 | 0.74 |
| 210 | 269 | 145 | 122 | 2 | 0 | 0.74 |
| 71 | 268 | 132 | 134 | 2 | 0 | 0.75 |
| 181 | 268 | 151 | 115 | 1 | 1 | 0.75 |
| 51 | 266 | 120 | 144 | 1 | 1 | 0.75 |
| 74 | 266 | 127 | 137 | 1 | 1 | 0.75 |
| 76 | 266 | 122 | 142 | 1 | 1 | 0.75 |
| 140 | 266 | 148 | 116 | 1 | 1 | 0.75 |
| 59 | 264 | 119 | 143 | 0 | 2 | 0.76 |
| 61 | 263 | 129 | 132 | 0 | 2 | 0.76 |
| 50 | 259 | 130 | 127 | 1 | 1 | 0.77 |
| 52 | 259 | 129 | 128 | 1 | 1 | 0.77 |
| 83 | 259 | 132 | 125 | 1 | 1 | 0.77 |
| 100 | 257 | 150 | 105 | 2 | 0 | 0.78 |
| 327 | 257 | 113 | 142 | 1 | 1 | 0.78 |
| 53 | 255 | 122 | 131 | 1 | 1 | 0.78 |
| 72 | 255 | 120 | 133 | 1 | 1 | 0.78 |
| 135 | 251 | 119 | 130 | 2 | 0 | 0.80 |
| 63 | 373 | 233 | 137 | 2 | 1 | 0.80 |
| 130 | 245 | 117 | 126 | 2 | 0 | 0.82 |
| 145 | 243 | 138 | 103 | 2 | 0 | 0.82 |
| 512 | 238 | 101 | 135 | 1 | 1 | 0.84 |
| 267 | 235 | 138 | 95 | 0 | 2 | 0.85 |
| 82 | 227 | 132 | 93 | 0 | 2 | 0.88 |
| 110 | 227 | 101 | 124 | 1 | 1 | 0.88 |
| 163 | 220 | 125 | 93 | 0 | 2 | 0.91 |
| 480 | 326 | 149 | 174 | 1 | 2 | 0.92 |
| 67 | 432 | 223 | 205 | 3 | 1 | 0.93 |
| 134 | 212 | 113 | 97 | 1 | 1 | 0.94 |
| 353 | 308 | 181 | 124 | 3 | 0 | 0.97 |
| 100 | 301 | 158 | 140 | 0 | 3 | 1.00 |
| 80 | 287 | 132 | 152 | 1 | 2 | 1.05 |
| 63 | 375 | 138 | 233 | 4 | 0 | 1.07 |
| 288 | 279 | 160 | 116 | 2 | 1 | 1.08 |
| 70 | 362 | 189 | 169 | 1 | 3 | 1.10 |

| | | | | | | |
|---|---|---|---|---|---|---|
| 81 | 267 | 149 | 115 | 3 | 0 | 1.12 |
| 52 | 266 | 169 | 94 | 3 | 0 | 1.13 |
| 180 | 262 | 127 | 132 | 3 | 0 | 1.15 |
| 175 | 261 | 119 | 139 | 0 | 3 | 1.15 |
| 79 | 334 | 145 | 185 | 1 | 3 | 1.20 |
| 77 | 330 | 214 | 112 | 0 | 4 | 1.21 |
| 91 | 246 | 141 | 102 | 0 | 3 | 1.22 |
| 79 | 324 | 152 | 168 | 0 | 4 | 1.23 |
| 71 | 238 | 129 | 106 | 3 | 0 | 1.26 |
| 75 | 288 | 164 | 120 | 2 | 2 | 1.39 |
| 104 | 284 | 142 | 138 | 0 | 4 | 1.41 |
| 81 | 282 | 130 | 148 | 1 | 3 | 1.42 |
| 145 | 274 | 91 | 179 | 4 | 0 | 1.46 |
| 566 | 255 | 80 | 171 | 3 | 1 | 1.57 |
| 65 | 310 | 181 | 124 | 1 | 4 | 1.61 |
| 168 | 370 | 218 | 146 | 6 | 0 | 1.62 |
| 52 | 354 | 146 | 202 | 6 | 0 | 1.69 |
| 452 | 294 | 162 | 127 | 0 | 5 | 1.70 |
| 59 | 345 | 119 | 220 | 0 | 6 | 1.74 |
| 338 | 339 | 210 | 123 | 5 | 1 | 1.77 |
| 85 | 323 | 151 | 166 | 6 | 0 | 1.86 |
| 51 | 266 | 140 | 121 | 0 | 5 | 1.88 |
| 115 | 260 | 137 | 118 | 4 | 1 | 1.92 |
| 107 | 306 | 123 | 177 | 0 | 6 | 1.96 |
| 103 | 305 | 135 | 164 | 1 | 5 | 1.97 |
| 168 | 295 | 187 | 102 | 3 | 3 | 2.03 |
| 71 | 240 | 107 | 128 | 4 | 1 | 2.08 |
| 84 | 324 | 164 | 153 | 7 | 0 | 2.16 |
| 101 | 267 | 116 | 145 | 0 | 6 | 2.25 |
| 68 | 264 | 114 | 144 | 2 | 4 | 2.27 |
| 50 | 292 | 113 | 172 | 0 | 7 | 2.40 |
| 91 | 249 | 101 | 142 | 0 | 6 | 2.41 |
| 550 | 247 | 126 | 115 | 6 | 0 | 2.43 |
| 59 | 201 | 100 | 96 | 0 | 5 | 2.49 |
| 93 | 352 | 184 | 159 | 9 | 0 | 2.56 |
| 87 | 265 | 133 | 125 | 2 | 5 | 2.64 |
| 98 | 324 | 189 | 126 | 8 | 1 | 2.78 |
| 118 | 287 | 144 | 135 | 6 | 2 | 2.79 |
| 123 | 356 | 210 | 136 | 9 | 1 | 2.81 |
| 52 | 239 | 104 | 128 | 0 | 7 | 2.93 |
| 74 | 302 | 143 | 150 | 8 | 1 | 2.98 |
| 67 | 267 | 143 | 116 | 0 | 8 | 3.00 |
| 83 | 360 | 161 | 188 | 9 | 2 | 3.06 |
| 66 | 276 | 143 | 124 | 0 | 9 | 3.26 |

| | | | | | | |
|---|---|---|---|---|---|---|
| 68 | 296 | 193 | 93 | 10 | 0 | 3.38 |
| 114 | 353 | 149 | 192 | 1 | 11 | 3.40 |
| 193 | 346 | 165 | 169 | 12 | 0 | 3.47 |
| 71 | 258 | 106 | 143 | 3 | 6 | 3.49 |
| 71 | 257 | 142 | 106 | 3 | 6 | 3.50 |
| 82 | 302 | 144 | 147 | 10 | 1 | 3.64 |
| 75 | 245 | 100 | 136 | 9 | 0 | 3.67 |
| 65 | 351 | 216 | 122 | 0 | 13 | 3.70 |
| 194 | 347 | 170 | 164 | 13 | 0 | 3.75 |
| 119 | 341 | 153 | 175 | 10 | 3 | 3.81 |
| 74 | 336 | 140 | 183 | 13 | 0 | 3.87 |
| 111 | 331 | 140 | 178 | 2 | 11 | 3.93 |
| 56 | 243 | 138 | 95 | 9 | 1 | 4.12 |
| 180 | 331 | 168 | 149 | 1 | 13 | 4.23 |
| 90 | 268 | 143 | 112 | 0 | 13 | 4.85 |
| 57 | 274 | 120 | 140 | 9 | 5 | 5.11 |
| 57 | 273 | 136 | 123 | 8 | 6 | 5.13 |
| 86 | 367 | 203 | 145 | 19 | 0 | 5.18 |
| 1442 | 309 | 196 | 97 | 15 | 1 | 5.18 |
| 94 | 307 | 174 | 117 | 5 | 11 | 5.21 |
| 54 | 320 | 189 | 114 | 0 | 17 | 5.31 |
| 73 | 446 | 160 | 261 | 14 | 11 | 5.61 |
| 127 | 331 | 125 | 187 | 19 | 0 | 5.74 |
| 84 | 299 | 174 | 107 | 15 | 3 | 6.02 |
| 190 | 328 | 169 | 139 | 4 | 16 | 6.10 |
| 71 | 426 | 140 | 259 | 27 | 0 | 6.34 |
| 50 | 266 | 116 | 133 | 16 | 1 | 6.39 |
| 51 | 265 | 108 | 140 | 1 | 16 | 6.42 |
| 66 | 385 | 151 | 209 | 20 | 5 | 6.49 |
| 461 | 173 | 104 | 57 | 4 | 8 | 6.94 |
| 349 | 367 | 202 | 136 | 28 | 1 | 7.90 |
| 129 | 293 | 155 | 113 | 25 | 0 | 8.53 |
| 213 | 253 | 119 | 111 | 23 | 0 | 9.09 |
| 64 | 398 | 144 | 216 | 0 | 38 | 9.55 |
| 468 | 257 | 113 | 119 | 1 | 24 | 9.73 |
| 86 | 234 | 126 | 84 | 0 | 24 | 10.26 |
| 112 | 258 | 125 | 100 | 33 | 0 | 12.79 |
| 115 | 348 | 137 | 165 | 46 | 0 | 13.22 |
| 62 | 403 | 179 | 168 | 52 | 4 | 13.90 |
| 192 | 417 | 200 | 159 | 58 | 0 | 13.91 |
| 50 | 285 | 133 | 112 | 39 | 1 | 14.04 |
| 55 | 314 | 106 | 162 | 0 | 46 | 14.65 |
| 165 | 323 | 117 | 158 | 1 | 47 | 14.86 |
| 473 | 261 | 78 | 140 | 0 | 43 | 16.48 |

| | | | | | | |
|---|---|---|---|---|---|---|
| 257 | 357 | 181 | 111 | 65 | 0 | 18.21 |
| 948 | 322 | 128 | 131 | 0 | 63 | 19.57 |
| 202 | 337 | 129 | 102 | 40 | 66 | 31.45 |

Table S5. Additional details of studies used in the summary of recombination dynamics. To the extent possible, we limited comparisons to repeats ≥ 50 bp with ≥ 80% identity and imposed a minimum isomer relative abundance of 1.6% to accommodate varying depths of sequencing among studies. Values were extracted from mostly from supplementary tables or precisely derived from vectorized figures using Inkscape 0.91. If only clone count was presented for alternative isomers, we used the average sequencing depth to estimate their frequencies. Citations are given in the main text.

| Species | # repeats ≥ 1 kbp analyzed | # repeats ≥ 1 kbp with ACGs ≥ 1.6% | # repeats < 1kbp analyzed | # repeats < 1 kbp with ACGs ≥ 1.6% | Notes |
|---|---|---|---|---|---|
| *Chrysanthemum nankingense* | na | na | 12 | 2 | Minimum identity of 95% |
| *Daucus carota* | 4 | 1 | 55 | 0 | We consider the 35 kbp overlapping repeat as a single large repeat. Recombination estimate of large repeats is from Southern hybridization, small from Illumina paired-end reads. Minimum identity of 90% |
| *Eucalyptus grandis* | 5 | 2 | 25 | 1 | AC frequency calculate using the mean (700x) coverage. Minimum identity of > 95% and length > 100 bp |
| *Monosonia ciliata* | - | - | 5 | 0 | Minimum repeat length of 100 bp |
| *Monsonia herrei* | na | na | 13 | 0 | Only repeats >100 bp and <800 bp were analyzed. |
| *Welwitschia mirabilis* | na | | 95 | 0 | Minimum length of 100 bp |
| *Nymphaea colorata* | 2,224 | 2 | 884,759 | 0 | Overlapping repeats |
| *Psilotum nudum* | 3 | 0 | 120 | 4 | Analyzed repeats between 100 bp and 5 kbp; minimum distance between repeats is 10kbp. Repeat of 1.1 kbp could not be analyzed due to the proximity of the two copies. |
| *Silene noctiflora* | 76 | 59 | 432 | 0 | Minimum identity of 95% |
| *Ophioglossum californicum* | 2 | 1 | 31 | 0 | Analyzed repeats between 100 bp and 5 kbp; minimum distance between repeats is 10 kbp |
| *Silene vulgaris* | 1 | 1 | 49 | 7 | Minimum identity of 95% |
| *Silene conica* | 42 | 20 | 690 | 42 | 5% of repeats < 200 bp were randomly sampled. Minimum identity of 95% |

| | | | | | |
|---|---|---|---|---|---|
| *Ginkgo biloba* | 3 | 3 | 31 | 0 | Minimum length of 100 bp |
| *Cucumis sativus* | 3 | 3 | 137 | 11 | A subset of repeats > 100bp were assessed for recombination. |
| *Picea abies* | 2 | 1 | 596 | 76 | |
| *Vigna angularis* | 1 | 1 | 43 | 11 | |
| *Mimulus guttatus* | 3 | 3 | 340 | 128 | Minimum repeat size is 40 bp, with a maximum of two differences. Minimum detection frequency is ~4%, assuming an average depth of coverage of 25x |
| *Viscum scurruloideum* | - | - | 35 | 23 | Only repeats ≤ 100 bp were analyzed |

Table S6. Estimates of the absolute number of mitogenome rearrangements and rearrangement rates in five species pairs with similar levels of divergence

| Species 1 | Species 2 | Divergence time | DNA shared (%) | Number of re-arrangements | Rearrangements per MY |
|---|---|---|---|---|---|
| *Vigna radiata* | *Glycine max* | 19 | 53.5 | 55 | 1.45 |
| *Capsicum annuum* | *Nicotiana tabacum* | 24 | 50 | 67 | 1.40 |
| *Brassica napus* | *Arabidopsis thaliana* | 28 | 54.5 | 59 | 1.06 |
| *Citrullus lanatus* | *Cucurbita pepo* | 30 | 43 | 67 | 1.12 |
| *Picea abies* | *Picea glauca* | 15 | 54.5 | 1,292 | 43.07 |

Table S7. Relative synonymous (dS) and nonsynonymous (dN) substitution rates inferred by the maximum-likelihood method implemented in PAML.

| Branch | dS | dN |
|---|---|---|
| *Pinus sylvestris* | 0.001867 | 0.000828 |
| *Pinus taeda* | 0.000831 | 0.000494 |
| *Pinus sylvestris-taeda* | 0.009509 | 0.005492 |
| *Pinus strobus* | 0.005668 | 0.004051 |
| *Pinus* spp. | 0.002182 | 0.000351 |
| *Picea abies* | 0.007241 | 0.003612 |
| *Picea glauca* | 0.001021 | 0.0012 |
| *Picea* spp. | 0.002279 | 0.001226 |
| Pinaceae | 0.014302 | 0.008863 |
| *Welwitschia mirabilis* | 0.132726 | 0.078619 |
| *Gnetum gnetum* | 0.046484 | 0.01966 |
| Gnetophytes | 0.073173 | 0.033236 |
| Gnetophytes-Pinaceae | 0.004826 | 0.003379 |
| *Araucaria heterophylla* | 0.179607 | 0.068664 |
| Pinales | 0.04667 | 0.009809 |
| *Ginkgo* | 0.038105 | 0.009994 |
| Ginkgo-Pinales | 0.022009 | 0.006466 |
| *Cycas taitungensis* | 0.036201 | 0.011118 |
| Gymnosperms | 0.129797 | 0.032194 |
| *Amborella tricopoda* | 0.002701 | 0.014727 |

Table S8. Organelle genomes and accession numbers used for support vector machine (SVM) training.

| Species | Genome | Accession |
|---|---|---|
| *Abies sibirica* | Plastid | Lars Arvestad, unpublished |
| *Amborella trichopoda* | Mito | KF754803.1 |
| | | KF754802.1 |
| | | KF754801.1 |
| | | KF754800.1 |
| | | KF754799.1 |
| | Plastid | NC_005086.1 |
| *Arabidopsis thaliana* | Mito | NC_001284.2 |
| | Plastid | NC_000932.1 |
| | Nuclear | NC_003070.9 |
| | | NC_003071.7 |
| | | NC_003074.8 |
| | | NC_003075.7 |
| | | NC_003076.8 |
| *Cycas taitungensis* | Mito | NC_010303.1 |
| | Plastid | NC_009618.1 |
| *Gnetum gnemon* | Plastid | Lars Arvestad, unpublished |
| *Juniperus communis* | Plastid | Lars Arvestad, unpublished |
| *Picea abies* | Plastid | NC_021456.1 |
| *Pinus sylvestris* | Plastid | Lars Arvestad, unpublished |
| *Populus trichocarpa* | Mito | KM091932.1 |
| | Plastid | NC_009143.1 |
| | Nuclear | Phytozome v. 10.0 |
| *Ricinus communis* | Mito | NC_015141.1 |
| | Plastid | NC_016736.1 |
| *Silene vulgaris* | Mito | NC_016406.1 |
| | | NC_016170.1 |
| | | NC_016402.1 |
| | | NC_016415.1 |
| | Plastid | NC_016727 |
| *Spirodela polyrhiza* | Mito | NC_017840.1 |
| | Plastid | NC_015891.1 |
| *Taxus baccata* | Plastid | Lars Arvestad, unpublished |
| *Triticum aestivum* | Mito | GU985444.1 |
| | Plastid | NC_002762 |

Figure S1. Frequency of direct and inverted repeats in the *Picea abies* mitogenome by length (log scale).

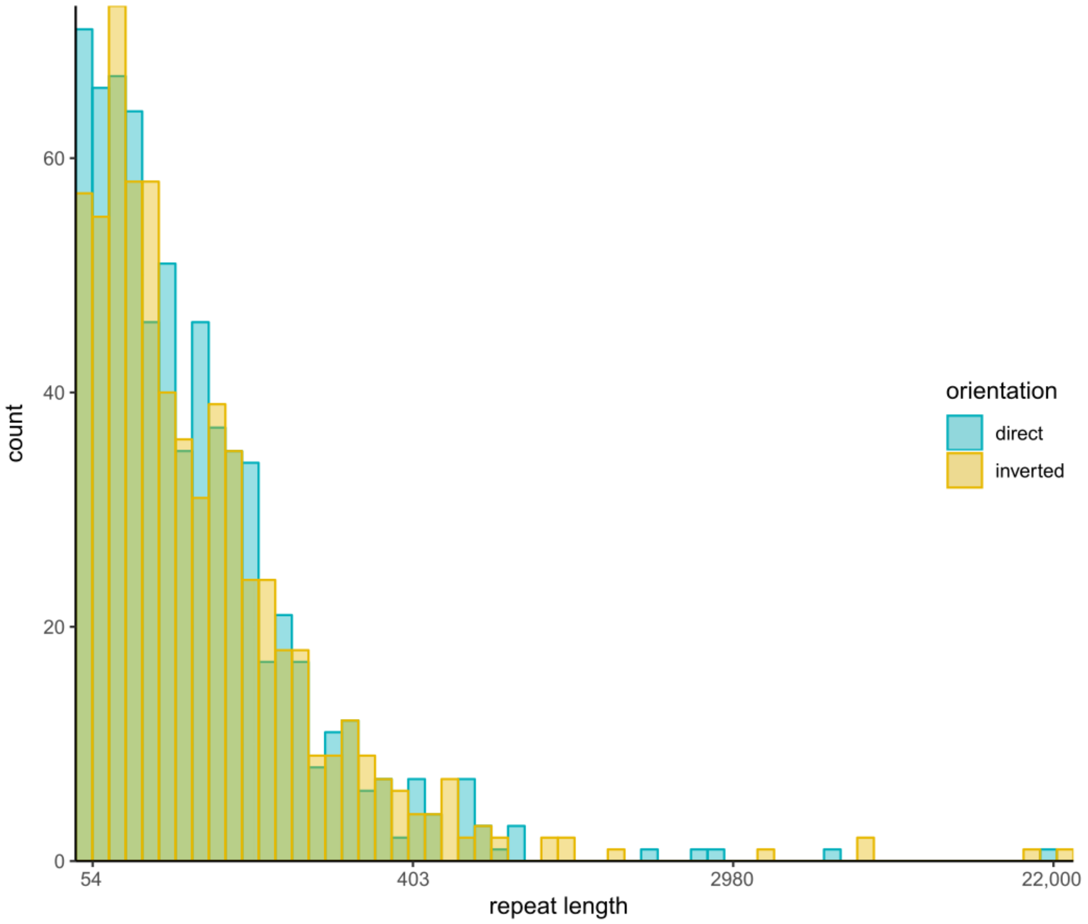

Figure S2. Schematic representation of the detection of alternate genome configurations produced by intramolecular recombination of inverted repeats. Colored boxes represent single-copy regions flanking the repeat, shown in white. Black lines under the boxes depict high quality, primary alignments of long reads that span the repeat and both single copy regions. Repeat Copy 1 and 2 are captured by the reference mitogenome assembly and thus are expected to be the most abundant configurations. The sequences for Alternative Genome Configurations 1 and 2 are the expected products of recombination between Repeat Copy 1 and Repeat Copy 2. In this case, long reads map to all four genome configurations and the two reference states are equally abundant and more common than the alternative genome configurations. The alternative genome configurations together comprise 33% of the read pool.

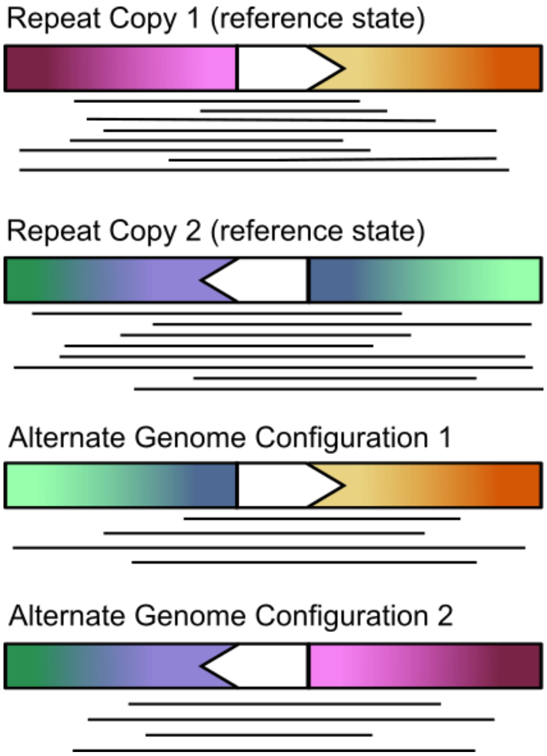

Supplementary Methods 1. Isolation of crude mitochondria from Norway spruce buds for mtDNA isolation

Keep all the equipment and buffers at 4°C before the isolation. Conduct all the steps on ice or at 4°C.

1. Thaw the buds (in storage medium - 10 mM MOPS, 5% DMSO, 5% glycerol, pH 7.3) in falcon tube) in a beaker with warm water. Change the water whenever it gets cold and invert the samples regularly to check if there is still ice and mix the solution. Keep doing until no ice is still visible and the leaves are soft
2. Pour the storage medium out of the falcon tube and dry the buds with adsorbent paper.
3. Weigh the material, dividing it into more samples with equal amount (max 10 g each)
4. Homogenize the tissue with a blender (keep the blender sloping giving short burst 2-3 s, about 10, and a final one of 5-10 s) in ~50 ml of grinding medium (see below - remember to add PVPP and PVP-40) and grind it with mortar and pestle (g fresh weight of tissues/ml of grinding medium: 1/5.
5. Filter the extract through a double layer of cheesecloth and a 100 µm mesh nylon net. Collect in a 20 ml centrifuge tube with screw lid.
6. Centrifuge at 2500 g x 5 min, 4 °C (Pellets containig nuclei, cell walls, starch, intact plastids).
7. Transfer the supernatant into new tubes (20 ml) by decanting without transferring the pellets. While decanting remember to slope the tube on the side where the pellet is.
8. Centrifuge supernatant at 13000 g x 15 min, 4°C (Pellets mitochondria, peroxisomes, plastids, fragments) in 1.5 ml Eppendorf tubes, before discarding it. Samples stored at -20°C .
9. The mitochondrial pellet is resuspended in 7 ml DNAse-I buffer (see below). Remember to add the buffer gradually pipetting gently in order to resuspend all the pellet. Do not add all the buffer at once. Use this step to collect all the pellets from one sample into one tube.
10. Dissolve 10.000 units of DNase I recombinant, grade I (Roche, Mannheim, Germany) in 1 ml DNAse buffer and add 10 µl to the mitochondrial suspension.
11. Incubate room temperature for 1 h to degrade any nuclear and chloroplast DNA present outside the mitochondria.
12. Terminate the digestion by adding 0.5 M EDTA (pH 8.0) to a final concentration of 25 mM (350 µl) and heat at 70°C for 15 min
13. Re-pellet the mitochondria by centrifuging at 16,000 g for 10 min
14. Discard the supernatant and resuspend the pellet in one tube with 20 ml wash buffer.
15. Wash once by re-pelleting at 16,000 g for 10 min, 4°C and discard the supernatant.

16. Resuspend in 360 µl of digestion solution using the p1000 with filter tips (you can freeze overnight if necessary or leave 2 h on ice to get rid of the foam). Add 40 µl of proteinase K solution and mix by vortexing.
17. Proceed with the protocol for Gram-Negative Bacteria Genomic DNA Purification Protocol (Thermoscientific).

Grinding buffer: 0.35 M mannitol, 30 mM MOPS, 2.5 mM $MgCl_2$, 2 mM sodium metabisulfite, 3 mM cysteine, 3 mM dithiothreitol, 1.25 mM EGTA, pH 7.4. Add 0.3% (w/v) PVPP and 0.3% PVP-40 (w/v) just before use.

DNAse buffer: 0.5 M mannitol, 10 mM $MgCl_2$, 50 mM Tris-HCl, pH 8, NO EDTA
